# Shaping Synthetic Multicellular and Complex Multimaterial Tissues via Embedded Extrusion-Volumetric Printing of Microgels

**DOI:** 10.1101/2023.05.17.541111

**Authors:** Davide Ribezzi, Marième Gueye, Sammy Florczak, Franziska Dusi, Dieuwke de Vos, Francesca Manente, Andreas Hierholzer, Martin Fussenegger, Massimiliano Caiazzo, Torsten Blunk, Jos Malda, Riccardo Levato

## Abstract

In living tissues, cells express their functions following complex signals from their surrounding microenvironment. Capturing both hierarchical architectures at the micro- and macroscale, and anisotropic cell patterning remains a major challenge in bioprinting, and therefore a bottleneck towards creating physiologically relevant models. Addressing this limitation, we introduced a novel technique, termed Embedded Extrusion-Volumetric Printing (EmVP), converging extrusion-bioprinting and layer-less, ultra-fast volumetric bioprinting, allowing to spatially pattern multiple inks/cell types. Light-responsive microgels were developed as permissive microenvironment for cell homing and self-organization, and as bioresins (µResins) for light-based bioprinting. Tuning the mechanical and optical properties of these gelatin-based microparticles enables their use as support bath for suspended extrusion printing, in which features containing high cell densities can be easily introduced. µResins can then be sculpted within seconds with tomographic light projections into centimetre-scale, granular hydrogel-based, convoluted constructs. Interstitial microvoids within microgels enhanced differentiation of multiple stem/progenitor cells (vascular, mesenchymal, neural), otherwise not possible with conventional bulk hydrogels. As proof-of-concept, EmVP was applied to create complex synthetic biology-inspired intercellular communication models, where adipocyte differentiation is regulated by optogenetic-engineered pancreatic cells. Overall, EmVP offers new avenues for producing regenerative grafts with enhanced functionality, and for developing engineered living systems and (metabolic) disease models.

## 1. INTRODUCTION

Biofabrication technologies are powerful tools to shape the architecture and guide the functionality of complex tissue constructs, and promise to revolutionize biomedical research and regenerative medicine. Through bioprinting and bioassembly techniques, precise control over the spatial organization of multiple cell types and biomaterials is possible via automated three-dimensional (3D) patterning processes.[1,2] Major challenges in bioprinting of functional constructs with native like-behavior, still rely on the use of biomaterials that provide a permissive environment for cell-driven self-organization and intercellular communication. While this is generally possible with hydrogels with low elastic moduli, ensuring high printing resolution and shape fidelity of these materials remains a key bottleneck, especially when using conventional layer-by-layer fabrication techniques.[3] Volumetric bioprinting (VBP) was recently introduced to increase the printing speed and overcome the geometric constraints of conventional layer-wise bioprinting approaches, even when using hydrogels with low mechanical properties.[4,5] Briefly, inspired by computed tomography, VBP relies on the projection of a series of 2D patterned optical light fields sent from different angles within a volume of a photopolymer, typically a cell-laden light-responsive hydrogel, also termed bioresin.[6–8] The cumulation of the light patterns form an anisotropic 3D dose distribution that triggers the polymerization of the bioresin only in the regions corresponding to the desired object.[4] However, the possibility to create high resolution features comprising multiple, independent structural elements intertwined into a construct remains a major challenge, especially when using multiple materials and cell types in a single printing process. Moreover, given that this technique is still in its infancy, there is still a limited array of available, printable materials, the vast majority of which do not provide environments in which cells can easily migrate and reorganize (or differentiate) into structures that mimic the native environment of human tissues. Enabling volumetric bioprinting using microgel-based materials, can offer a new versatile solution to these limitations, allowing to create complex 3D structures embedding features containing multiple inks, multiple cell types, in which bioprinted cells can rapidly grow and home.

Microgel-based biomaterials, also known as granular hydrogels, are an emerging class of biomaterials and a promising alternative to conventional hydrogels for 3D bioprinting applications.[9] Granular hydrogels consist in packed assemblies of hydrogel microparticles (the microgels), typically produced using methods, such as mechanical fragmentation of bulk gels,[10,11] via microfluidic systems,[12,13] lithography[14,15] or batch emulsions.[16,17] Microgels packing to obtain granular hydrogels can be performed by centrifugation,[18] gravity-assisted sedimentation,[19] or vacuum-driven removal of the liquid phase from a microgel suspension.[17,19–21] Notably, while individually each microgel possesses comparable properties to its bulk hydrogel counterpart, packed microgel assemblies can be designed and customized to display a broad array of macro-scale behaviors, for instance by tuning the properties of each individual microgel, or mixing microgels with different physico-chemical profile (*e.g.*, size, shape, mechanics, degradability).[22] Importantly, packed microgels produce micrometre-sized voids of interstitial space,[23] where cells can easily infiltrate, proliferate and spread without needing to degrade the hydrogel. Obviously, this further increases the degrees of freedom achievable in terms of biological performance and range of targetable biomedical applications.[18,19,24] Of particular relevance for the field of biofabrication, granular hydrogels have already made their way in techniques like extrusion-based printing as well, due to their unique rheological properties.[22] In fact, while on the one hand jammed microgels can be directly used as bioinks for extrusion bioprinting,[9,25,26] microgel slurries are also an ideal support bath for gel-in-gel printing, a process also known as embedded extrusion printing.[27–29] In conventional extrusion printing, many printing artefacts are caused by the deformation of extruded filaments due to gravity and surface tension-driven effects.[30] To overcome this problem, embedded extrusion printing leverages printing inside of a support bath, capable of keeping the printed structure in its intended 3D geometry and preventing it from collapsing, even when the extruded ink is a hydrogel with low viscosity and limited mechanical properties. When the support bath during the extrusion printing process is composed of a microgel slurry, the microgels around the moving nozzle locally displace, allowing the extrusion of the ink, and keeping the printed filament in place.[31] However, as for all extrusion techniques, the printing time increases cubically with the scaling factor, resulting in unfavourable printing times when cell-laden, centimeter-scale constructs are required.[4]

In this work, we demonstrate for the first time the convergence of VBP with embedded extrusion bioprinting, hereby termed <u>Em</u>bedded Extrusion <u>V</u>olumetric <u>P</u>rinting (EmVP), to rapidly produce multi-material, centimeter-scale microgel based-constructs, in which multiple cell types can be patterned at high cellular densities, typical of many native dense and soft tissues (*i.e*., pancreas, liver, etc.). With this approach, a key challenge of classical volumetric printing, that is the inherent difficulty in producing structures containing multi-ink and multi-cell features can be overcome. The workflow for EmVP described in this work, and the different steps to characterize the process and the materials utilized is summarized in **Figure 1**. As platform material for EmVP, we developed and characterized a photo-crosslinkable micro-resin (µResin) composed by jammed gelatin methacryloyl (gelMA) microgels (**Figure 1A**). These can be annealed in a spatial-selective fashion via volumetric bioprinting, to produce well-defined, microporous objects (**Figure 1B**), and the potential of the interstitial spaces between volumetrically annealed, printed microgels to support the culture of multiple cell types was investigated (**Figure 1C**). Notably, prior to the tomographic printing step, the µResin can be utilized as cell-laden support medium for embedded printing, leveraging on its particulate composition and its shear-thinning behavior, to pattern any cell type or material in precisely selected patterns within the bioresin volume (**Figure 1D**). This approach is particularly important both to recreate high cell density areas within volumetrically bioprinted constructs (**Figure 1E**), and also to minimize the number of cells needed during the subsequent volumetric print. Moreover, we also showed the possibility for including mechanically strong inks to provide further structural stability to the final constructs. Finally, as proof-of-concept of application, we introduced a new tissue engineering approach, in which we use synthetic biology tools and optogenetics to guide the communication between different bioprinted cell types (**Figure 1F**). Here, EmVP is employed to produce a spatially defined, co-culture system, in which a precisely patterned pancreatic β-like cell line, engineered to release insulin upon blue light exposure, interacts with a stromal tissue composed of (pre-)adipocytes, in the perspective of developing an *in vitro* synthetic model that can have future applications *e.g.,* in studying metabolic dysregulation in obesity and diabetes.

**Figure 1:**
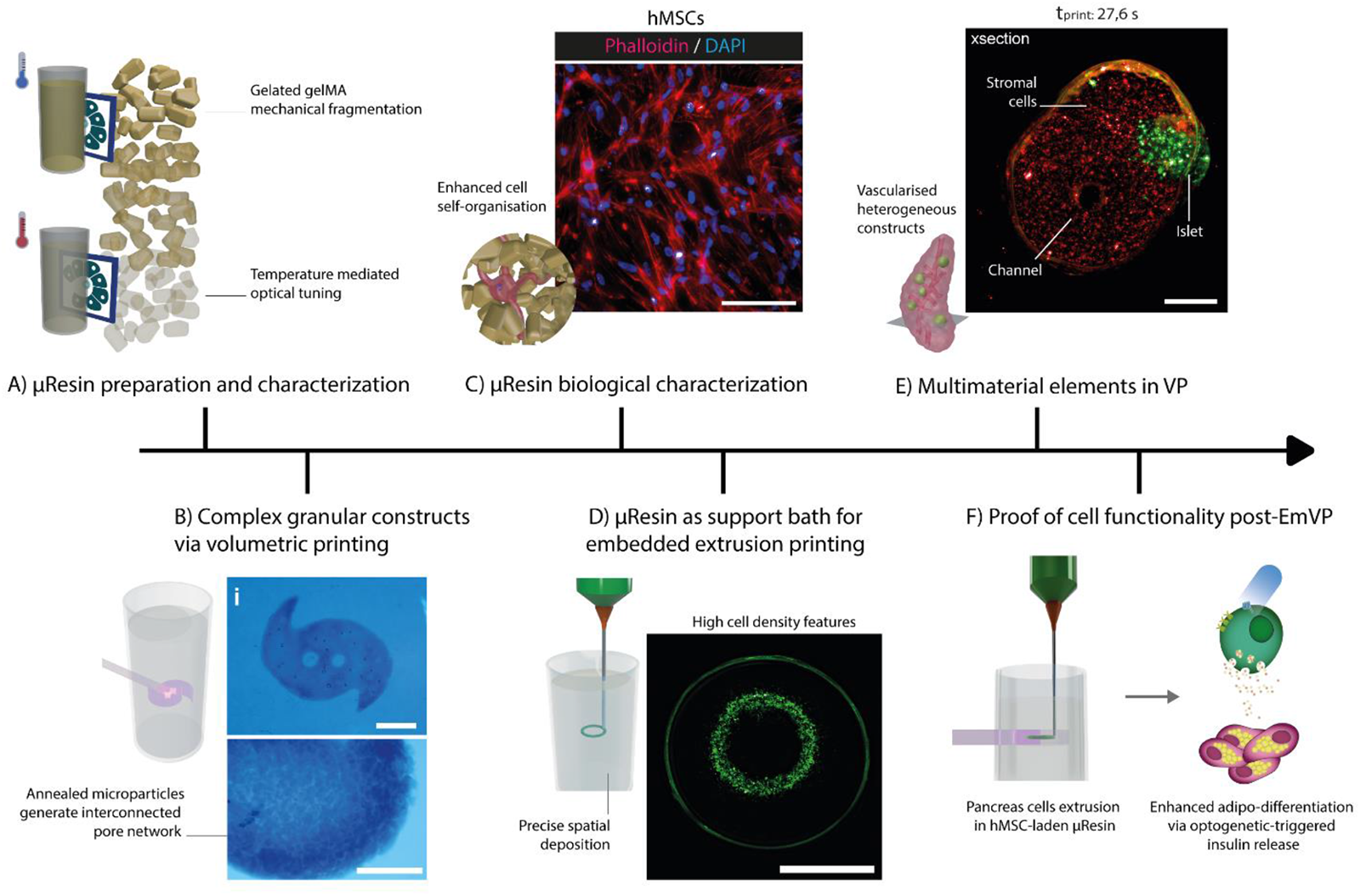
Workflow detailing the different steps towards the development and characterization of the Embedded Extrusion Volumetric Printing process, as a new biofabrication approach to enable microgel constructs and multimaterial elements in volumetric printing. A) Preparation and characterization of a µResin composed of gelatin-derived microparticles. Mechanical, rheological and optical properties are analyzed and optimized to ensure biological performance and transparency suitable for the tomographic printing process. B) The µResin were proven to be a suitable biomaterial to create granular structures via volumetric printing. Scale bars: 1 mm. C) Biological characterization of the µResin, which offers a soft and highly porous microenvironment to the encapsulated cells. Scale bar: 100 µm. D) The µResin bath was then studied for its ability to permit embedded extrusion bioprinting, allowing to produce localized regions and precisely patterned structures containing high cell concentration. Scale bar: 4 mm. E) These cell-dense areas become heterocellular element in more complex architectures upon performing the Volumetric (bio)Printing step, to impart the final 3D shape to multicellular and multimaterial constructs Scale bar = 1 mm. F) Finally, a proof-of-functionality of cells subjected to the whole EmVP process was provided, utilizing extruded iβ-pancreatic cells, secreting insulin upon blue light exposure, which influenced the adipogenic differentiation of hMSC loaded into the µResin.

## 2. RESULTS AND DISCUSSION

### 2.1 µResins for volumetric bioprinting with tunable optical and mechanical properties

To generate a microgel-based bioresin, we first focused on the technology to produce light-sensitive particles. Although microgel slurries produced from virtually any light-responsive hydrogel could be used for EmVP, gelMA was selected as proof-of-concept material for this study, due to its wide-spread use in bioprinting,[32] due to its biocompatibility and tunability for a large range of applications in both extrusion-based and light-based bioprinting.[4,33,34] In this work, microgels were produced by mechanically breaking a thermally gelated, non-photoexposed, gelMA bulk hydrogel into microscale particles using a simple rotational blender, allowing to turn bulk hydrogels into microparticles within seconds or few minutes.[11,22] Particles obtained in this way show an irregular, polygonal-like morphology (**Supplementary Figure S1**). Obtaining a high amount of microgel in a fast way is crucial when the granular hydrogel is meant to be used as a bioresin in a volumetric printing process to generate cm^3^-scale objects. We first assessed the effect of four different milling times (45s, 60s, 90s, and 120s) on microparticle size, geometry, and on the rheological profile of the microparticle slurries (**Figure 2A**). Results showed that blending time significantly affects the particle size after 45s, and the smallest dimension (159 ± 75 µm) was achieved after 90s. Irrespective of the blending time, all the µResin formulations showed a marked shear thinning behavior (**Figure 2B**), compatible with what is commonly reported for granular hydrogels,[22] and a key parameter for ideal materials intended not only as bioinks for extrusion printing,[35] but also as support baths for embedded bioprinting.[11] Next, the obtained slurries can be converted into a cohesive scaffold by photocrosslinking, resulting in the formation of both intra- and inter-particle covalent bonds, to create microporous annealed gels.[36] The photo-crosslinking was achieved by supplementing the µResin with 0.1% w/v Lithium phenyl(2,4,6-trimethylbenzoyl)phosphinate (LAP), a photo-initiator commonly used for biofabrication purposes due to its optimal water solubility and high cytocompatibility.[37,38] The mechanical properties of the photo-annealed constructs were then assessed while maintaining the gels at the same temperature (5°C) used during the microgel fabrication phase. Unconfined compression tests revealed a statistically significant correlation between the milling time and the compressive moduli, showing how a significant drop of the mechanical properties follows a reduction of the microgels dimension (from 14.3 ± 2.2 to 5.6 ± 0.9, for the samples 45s and 120s, respectively; p <= 0.05), which was always consistently lower than the values found for bulk hydrogels (46.3 ± 5.7). This decrease is consistent with the smaller size distribution, found for microgels produced upon longer blending time. High size polydispersity could in fact allow tighter packing, as smaller particles fill the interparticle voids between larger microgels, therefore, resulting in overall stiffer structures.[39] The stiffness of the gels is an important characteristic as it influences the overall mechanical properties, as well as the behavior and morphology of the incapsulated cells.[40] Notably, the mechanical performance of the annealed microgels can also be tuned adjusting the temperature at which the thermal gels are processed prior to photoexposure. Consistent with the literature, our results showed how photocrosslinking after the exposure to higher temperatures (specifically, 21°C, **Figure 2C**), led to softer gels (modulus values from 8.7 ± 0.8 to 3.2 ± 0.3, for the samples 45s and 120s, respectively; p <= 0.0001), compared to those obtained at lower temperatures, known to increase the amount of triple helices in the gelatin backbone.[22,41–43] Therefore, control over the pre-annealing temperature offers an additional strategy to expand the range of tissue culture applications for the µResin materials, closer to the requirements needed for mimicking soft tissues (e.g., for pancreas or liver). In addition, smaller particle size is also beneficial to reduce the printing resolution.[44] With this in mind, with a compressive modulus of 4 ± 0.3 kPa and with the smallest microgel dimensions (159 ± 75 µm) observed in our study, we decided to proceed with the formulation of the µResin resulting from the 90s milling. Even though the 120s milling formulation showed even lower mechanical properties (compressive modulus: 3.2 ± 0.3 kPa), this specific condition resulted in uncontrolled dissolution of gelated GelMA during blending, ending up with way less volume of granular hydrogel compared to the starting bulk volume. This could be attributed to the local temperature increment during the mechanical fragmentation, with the consequent heating and dissolution of the physically-gelated polymer.

**Figure 2:**
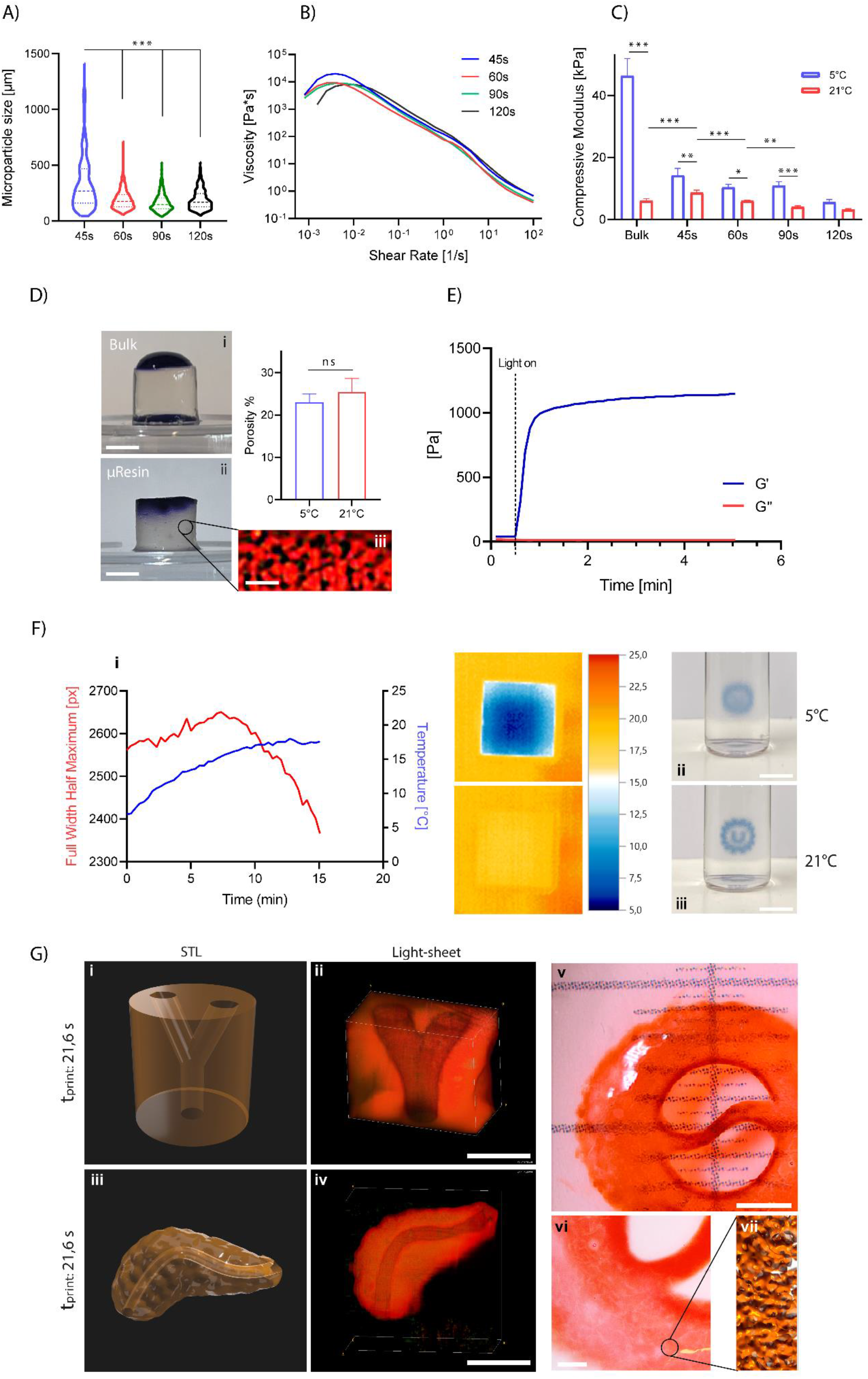
Characterisation of the photo-crosslinkable µResin based on jammed gelMA microparticles. A) quantification of the microparticles dimension related to the milling time of the thermally-gelated gelMA. B) Representative (n=3) rheological properties of the microgel slurries obtained at different milling times. C) Compressive mechanical properties of the microparticle-based gels (n=4) crosslinked at 5°C and 21°C. D) Porosity (n=6) characterization of (i) bulk and (ii) µResin gels, with a cross section of the (iii) µResin gel showing the particulate, porous microstructure. Scale bars: i,ii = 2.5 mm; iii = 200 µm. E) Representative (n=3) crosslinking kinetics after light exposure of the bioresin formulation obtained by milling for 90s. F) Characterisation of the optical properties (light scattering) of the microparticle-based bioresin as a function of the temperature (n=3), and micrographs paired with thermal photography of the vials containing the bioresin at (i) 5°C and (ii) 21°C. Scale bars: ii, iii = 6 mm. G) Volumetrically bioprinted models with additional negative features (open perfusable channels). Branched vasculature (i) CAD model and (ii) light sheet scan reconstructions, digitally cropped to the central region to facilitate 3D visualization of the hollow features. Pancreas models with duct, (iii) CAD model and (iv) light sheet scan reconstructions. Scale bars: 3 mm. Stereomicroscopy image of (vi) test print model: a hollow circular structure with a thin wavy central strut measuring obtained via volumetric printing of the bioresin formulation obtained after milling for 90s, with a (vi) close-up image to highlight the particulate nature of the constructs, and a (vii) 3D reconstruction of the particulate/porous network microstructure. Scale bars: ii, iv, v = 2 mm; vi = 500 µm.

To visualize and characterise the microstructure of the annealed gels, light-sheet microscopy was used to image interstitial pores. Cross-sections of the volumetric stacks (**Figure 2D**) revealed interconnected pores within the gels, with an overall porosity that did not significantly differ between gels annealed at 5°C and 21°C (showing 23 ± 2 % and 25 ± 3 % of pore fraction respectively). As the porosity is primarily dependent on inter-particle packing and packing density, this could be potentially modulated blending in the slurry particles produced with a different method, having different size, as previously demonstrated in other works on microgels.[20,22] The interconnected nature of the pore network, beneficial for nutrient diffusion and catabolite exchange during tissue culture, can also be qualitatively visualized by leaving a drop of stained solution onto the µResin based gel, which is rapidly percolated by dye, as opposed to what observed in bulk hydrogels.

Next, we investigated the potential of the selected microgel formulation, in terms of gelation kinetics, optical properties, and printing resolution. Showing a fast kinetic crosslinking and comparable to recently reported studies,[5,45,46] the µResin reached a storage modulus G’ plateau of 1,15 ± 0,072 kPa in 30 s after light exposure (**Figure 2E**). Despite of the promising fast gelation, however, the particulate nature of the µResin, in which microgels and void spaces alternate, is a primary cause of light scattering. As already observed,[5] scattered photons will distort the tomographic projections, altering the light dose distribution also in areas of the volume outside of the component that will be printed, therefore, resulting in printing artefacts. To address this limitation when printing with high scattering resins, our group previously demonstrated rescuing of high printing resolution (⁓40µm) via iodixanol-mediated optical tuning, a biocompatible compound to match the refracting index of highly scattering intracellular structures.[5] Alternatively, scattering effects can be compensated also via computational corrections of the tomographic light patterns delivered in the printing vial.[37] Here, we show how the optical properties of the µResins and, therefore, the scattering nature of the microgels, could be inherently modulated by tuning the temperature selected for the microgel slurries prior to the photocrosslinking-mediated annealing (**Figure 2F**). A custom-made optical setup was designed to evaluate the effect of light scattering as a function of the temperature. A laser beam (532 nm) having a circular cross-section was shined through the centre of a PMMA squared cuvette containing the µResin, and the resulting scattering pattern was projected on a photodetector placed behind the cuvette. For an ideal, non-scattering media, the photodetector should register a bright spot having the same size and homogeneous light intensity of the delivered laser, while a diffuse halo with a decreasing light intensity towards its outer border will be registered for scattering media. The full width half maximum (FWHM) of the patterns was then calculated from the center along the horizontal axes with a MATLAB script on different pictures taken at different time points as the µResin slowly warmed up to room temperature. A thermal camera was placed on top of the setup allowed to monitor the temperature of the µResin during the measurements, and the significant drop (**Figure 2Fi**) from 2561.5 ± 20.5 to 2381 ± 21.2 pixels of FWHM (p <= 0.01), respectively at 7°C and 18°C, clearly shows how higher pre-annealing temperatures can be exploited to reduce light scattering, likely due to lower triple helices content and therefore more amorphous nature of thermal gel,[41] and permit high resolution printing. To qualitatively visualize the magnitude of this effect, a drawing of logo of the University Medical Center Utrecht, printed on paper, was placed behind a printing vial containing the µResin, and pictures were taken with the µResin at ⁓5°C (temperature registered after putting the vial at 4°C, which is usually performed before the volumetric printing process;[4,5] **Figure 2Fii**) and ⁓ 21°C (temperature registered immediately after mixing the µResin with LAP; **Figure 2Fiii**), showing a dramatic improvement of the optical image for the latter, where the drawing was fully visible, as opposed to the fuzzy, undistinguishable pattern seen at ⁓ 5°C.

Having identified suitable conditions for microgel processing enabling the use as bioink or as support bath, we demonstrated the possibility to generate high-resolution volumetric constructs composed by annealed microgels via VBP. Large-scale constructs with positive and negative features (channels) were successfully printed at resolutions up to 244 ± 16 µm and 461 ± 26 µm for positive and negative features, respectively (**Figure 1B**), and an array of complex geometrical shapes and channel-laden hydrogels was produced in less than 22s (**Figure 2Gi/ii/iii/iv**). Notably, microscopic observation confirmed how the printed objects are composed of assembled microgels, and how these granular gels are inherently microporous even after the crosslinking (**Figure 2Gv**). Thus, exploiting the combination of interstitial microporosity and larger, printed hollow channels could eventually facilitate nutrient accessibility throughout the volume of bioprinted constructs during tissue culture. Finally, with regards to the printing resolution, previous reports on VBP demonstrated the ability of the technique to resolve features as small as 40 µm, when using bulk gelMA bioresins.[5] Differently from these homogenous resins, in line with the particulate nature of the microgel constructs, the minimum printing resolution when using µResins is instead limited by the particle size, and on how many particles are bound together by the tomographic light exposure in the proximity of small features, such as the spikes of the model in Figure 1B. Therefore, although our best resolution is in the range of approximately the size of two particles, in future studies further optimization of the volumetric printing resolution could be performed producing smaller microgels, for instance switching from gelatin blending to batch emulsion,[47] or coalescence-based strategies,[48] which, although requiring extensive purification steps to remove emulsifiers, or being lower throughput compared to our approach, showed minimum particle size in the range between 1 and 10 µm, and also enable improved resolution of extrusion-printed inks.

### 2.2 GelMA µResins provide a permissive environment for cell encapsulation and self-organization

Having identified suitable properties for volumetric printing, the biological performance of the annealed µResin was investigated (**Figure 3**). More specifically, the effects of the interstitial microporosity were analyzed by embedding multiple cell types in the µResin before photo-crosslinking. The granular gels proved to offer beneficial conditions to sustain the culture of multiple cell types, which typically thrive only in low stiffness (<1-2 kPa) or highly porous microenvironments. A common result in cell encapsulation in elastic, bulk hydrogels such as the gelMA used in this work (85% DoF, 5% w/v), is that cells tend to acquire a round morphology, and can remodel the hydrogel only slowly, through matrix degradation, effectively preventing direct cell-cell contact and interactions.[49] This same behavior was observed when culturing human bone marrow-derived mesenchymal stromal cells (hMSCs) in bulk hydrogel controls (**Figure 3A**), over 7 days of culture. Conversely, in the µResin, hMSCs displayed an elongated morphology, with a stretching area increased by more than 300%. Likewise, human umbilical vein endothelial cells (HUVECs) were able to grow, elongate, and connect forming pre-angiogenic capillary networks throughout the whole microgel-based constructs when co-cultured with hMSCs, showing a ⁓1800% increment on the number of junctions between the nascent capillaries, in contrast to what was observed in bulk GelMA gels (**Figure 3B**). This crucial characteristic makes granular gels promising candidates for tissue engineering purposes, including vasculature development.[50]

**Figure 3:**
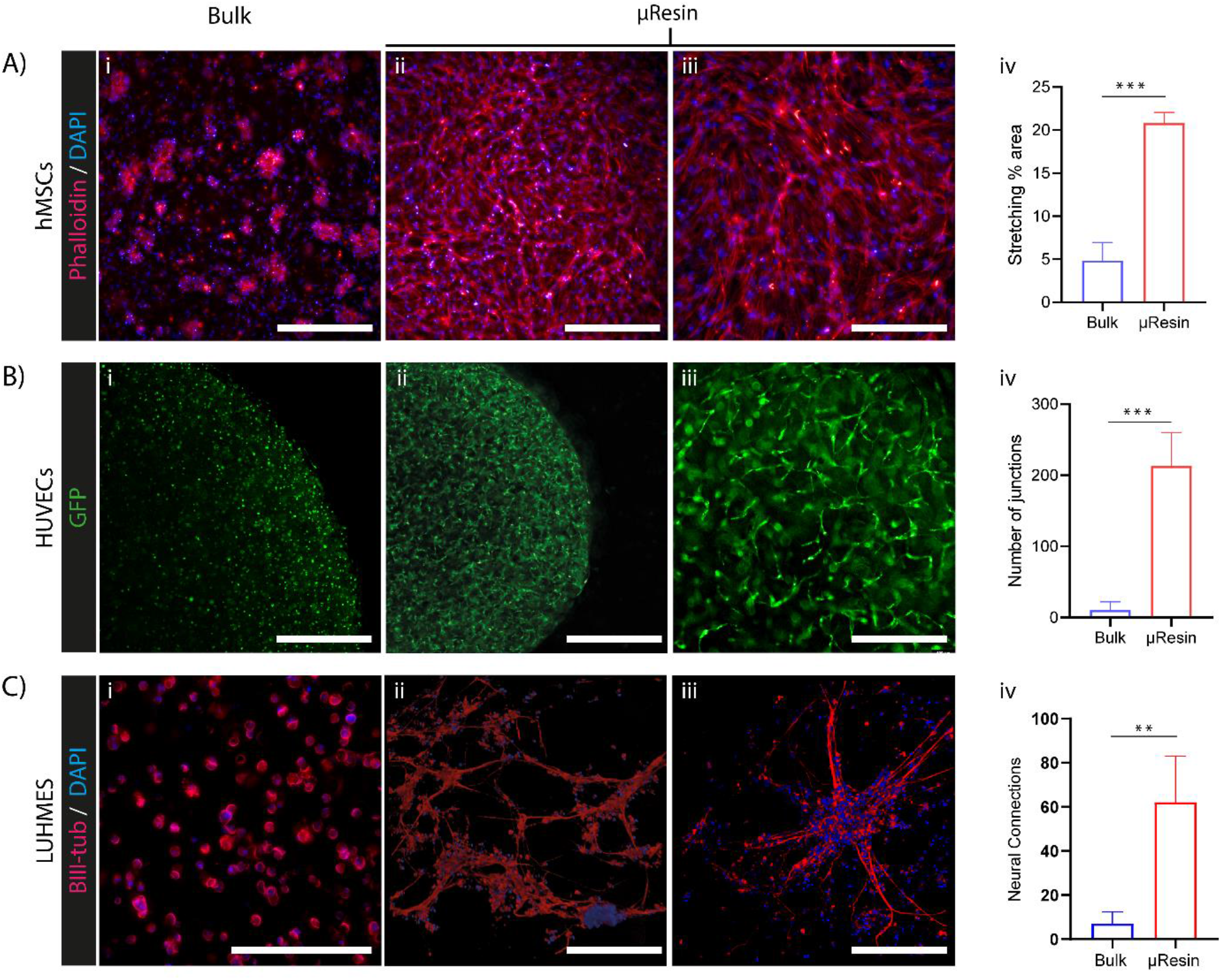
Effect of the microstructure of the microgel based gels on cell behavior. A) Morphology of hMSCs at a cell density of 5M cells/mL in (i) bulk (scale bar: 400 µm) (ii) gelMA µResin (scale bar: 400 µm), (iii) gelMA µResin (scale bar: 250 µm) and (iv) stretching % area quantification (day 7 post-encapsulation, n = 3). B) HUVEC cells at a density of 2.5M cells/mL in (i) bulk (scale bars: 750 µm) (ii) gelMA µResin (scale bar: 750 µm), (iii) gelMA µResin (scale bar: 400 µm) and (iv) quantification of the amount of nascent capillary junctions (day 7 post-encapsulation, n = 3). C) LUHMES cells at a cell density of 10M cells/mL in (i) bulk and (ii, iii) gelMA µResin (scale bars: i = 100 µm; ii,iii = 200 µm) and (iv) quantification of the amounts of neuronal junctions (day 8 post-encapsulation, n = 3).

As additional proof-of-concept of the suitability of our µResin for the recapitulation of soft tissues, Lund human mesencephalic neuronal cells (LUHMES) were embedded in the µResin to test neuronal differentiation, detecting the presence of the neuronal marker βIII-tubulin. Results showed how LUHMES in µResin can form neurites that are growing omnidirectionally in large numbers (**Figure 3C**). As these cells require basal membrane proteins as attachment points, this culture was enhanced with mixing into the µResin and the bulk gelMA low amounts of Matrigel (11.2 μl/ml), as well as fibronectin (50 μl/ml). These comprise ECM compounds typically used in neuronal cell culture,[51] and this experiments underlines the potential to easily functionalize the microgels with different matrix proteins. LUHMES were able to sprout and connect, forming neural connections throughout the whole µResin gels 785% more efficiently than in bulk gelMA, in which the cell-cell connections were observed only in the proximity of small cell clusters.

These results support the hypothesis that microporosity can be successfully leveraged to culture a broad array of cell types. Average length of the neurites changes as a function of local cell density, with protrusions measuring between ⁓50 μm to ⁓250 μm in densely populated regions of the µResin, and neurites spanning distances as long as ⁓700 μm when connecting more distant cells. Moreover, they confirm evidence from previous studies,[52] which employed microgels for human *in vitro* models of neural tissue, leading to improved cell-cell contacts and tissue maturation.[9,53,54] Further, it should be noted that, while this work focused on a single method for producing microgels, slurries obtained from particle-generation methods, [48] could be an useful future approach to further modulate inter-particle porosity, and therefore also cellular response.

### 2.3 µResins promote MSC adipogenic commitment even in insulin-depleted differentiation media

The specific properties of each microgel impact the macro-scale behavior of granular hydrogels, as well as the fate of the encapsulated cells. To further investigate how the stimuli coming from the architecture of the microenvironment can affect cell behavior, we embedded hMSCs in both the gelMA bulk hydrogel and the µResin and cultured them in adipogenic differentiation media (**Figure 4**). This choice is due to the overall compatible mechanical properties of the hydrogels, comparable with the stiffness of adipose tissue,[55,56] and for the central role of the adipose tissue in the endocrine regulation of our body, which is also implicated in key metabolic disfunctions, such as diabetes,[57,58] a disease of high interest for biomedical research.

**Figure 4:**
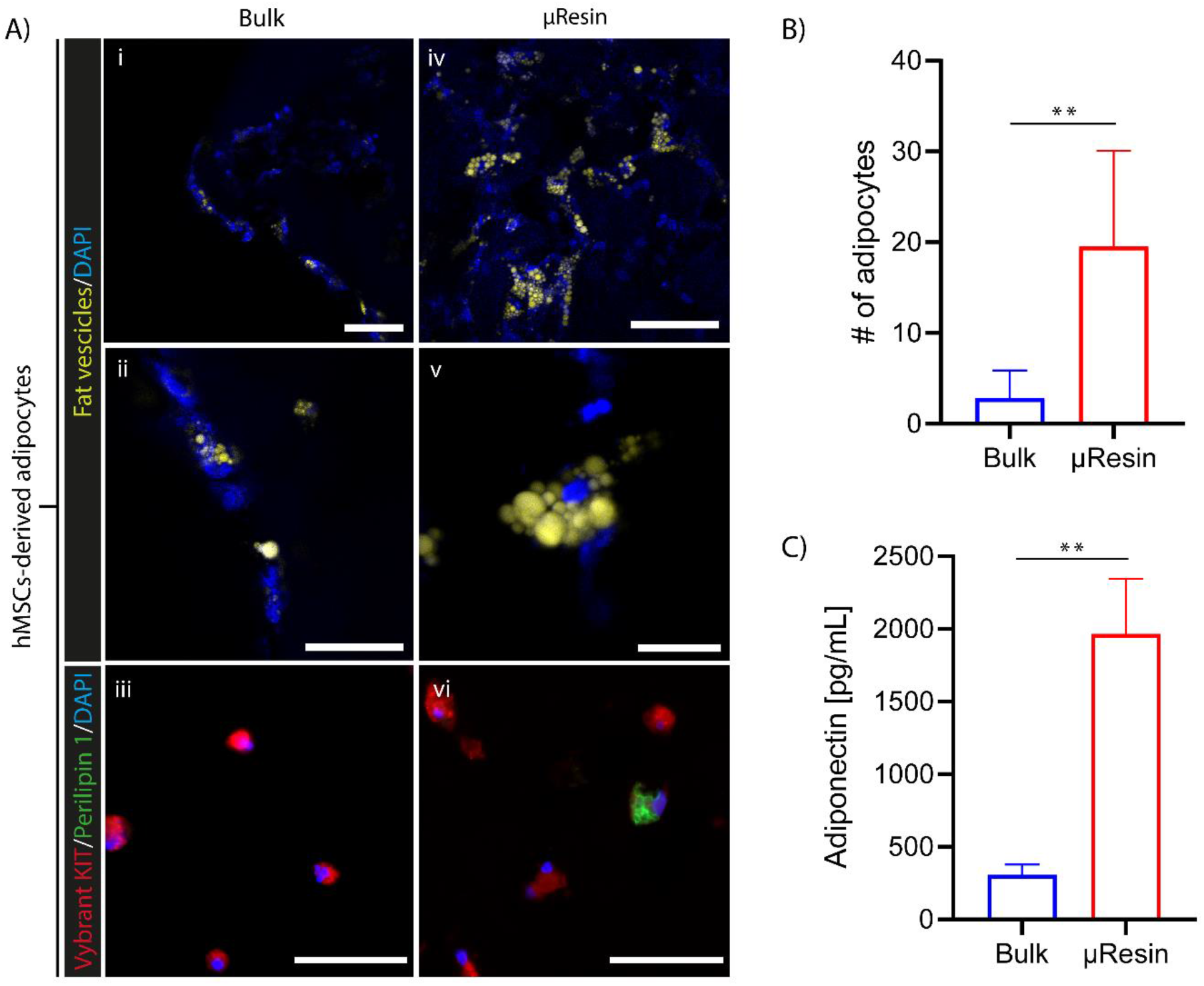
Effect of the microstructure of the hydrogels on hMSC cell differentiation into adipocytes. A) Comparison between gelMA bulk (i, ii, iii) and µResin (iv, v, vi) constructs, showing staining for the cell membrane (Vybrant DiD), fat vesicles (LipidTox) (whole mount staining) and Perilipin 1 immunostaining (cryosections). Nuclei are counterstained with DAPI. B) Quantification of the amount of adipocytes per field of view in bulk and gelMA µResin constructs, cultured without insulin in the differentiation medium (n=6). C) Adiponectin concentration in the supernatant of bulk and gelMA µResin constructs (n=3). Scale bars: i, ii, iv = 100 µm; iii, vi = 50 µm; v = 20 µm.

Interestingly, we demonstrated how the obtained microstructure is able to facilitate the adipogenic differentiation of hMSCs even in absence of insulin, a key factor often used in adipogenic media. In particular, a higher number of fat vesicles was observed (**Figure 4Ai/ii/iv/v**) and a significant higher percentage of differentiation was observed in the granular gels (20 ± 10 differentiated adipocytes per field view) compared to the bulk gels (3 ± 3 differentiated adipocytes per field view) (**Figure 4B**). Moreover, Perilipin 1 staining at the surface of lipid droplets was observed within differentiated MSC in µResin culture, whereas hardly any perilipin was detected in the bulk culture (**Figure 4Aiii/vi**). Finally, differentiated hMSCs in µResin culture showed markedly higher secretion of adiponectin (1965 ± 379.7 pg/mL), a major adipokine and hallmark of adipogenic differentiation, than in bulk culture (307.3 ± 70.1 pg/mL) (**Figure 4C**). All these results further confirmed how granular hydrogels address important cytocompatibility limitations of bulk encapsulation.

### 2.4 µResins can act as support bath for multi-material extrusion bioprinting

Next, as a necessary step towards enabling the generation of complex multimaterial and heterocellular constructs via combining embedded and volumetric printing, we assessed the characteristics of the µResin when applied as suspension bath. Embedded extrusion printing, or free-form three-dimensional (3D) printing, is a promising technique enabling researchers to produce complex shapes and overhangs, with a broad array of low-viscosity materials.[27,59] Notably, embedded printing has been used to print ink with high cell density (up to 1×10^7^),[60] and even inks composed solely of cell slurries.[61]

The shear-thinning behavior identified in our non-photocrosslinked microgel slurries (**Figure 1B**), typical of a non-Newtonian fluid, supported the notion that the µResin could be used as a suspension bath for embedded printing.[11,29] It is therefore possible to include specific pockets and structures made from multiple materials and cell types into the microgel constructs. To assess this, two extrudable (bio)inks were designed aiming to i) tune the mechanical bulk properties of the soft, annealed µResin, and ii) for precisely patterning high cell densities.

The first ink is based on a double network of gellan gum (GG) and poly(ethylene glycol) diacrylate (PEGDA), already proved to be a suitable blend for extrusion bioprinting, thanks to the shear thinning rheological profile of GG.[62] This material was investigated to provide mechanical reinforcement and to enhance the structural stability of printed objects. The second bioink was designed to allow for the delivery of high cell densities with good shape fidelity, even in absence of stably incorporated hydrogel carriers. For this purpose, we produced a bioink based on methylcellulose (MC), due to its ability to both modulate the viscosity of the cell suspension,[63–65] and rapidly dissolve post printing in an aqueous environment.

As reported in **Figure 5A**, both inks showed a decrease of the viscosity values when the shear rate increase, typical shear-thinning behavior crucial for a biomaterial to be employed as an extrudable bioink.[66] In particular, independently from the strain % applied, MC-based inks showed a higher loss modulus G” compared to the storage modulus G’, indicating how the viscoelastic properties are always dominant in this polymer at this specific concentration (2% w/v). In the GG/PEGDA blend, the properties of GG[67] were dominating the system making it shear thinning slightly above 10% of strain, where G” becomes greater than G’ (**Figure 5B**).

**Figure 5:**
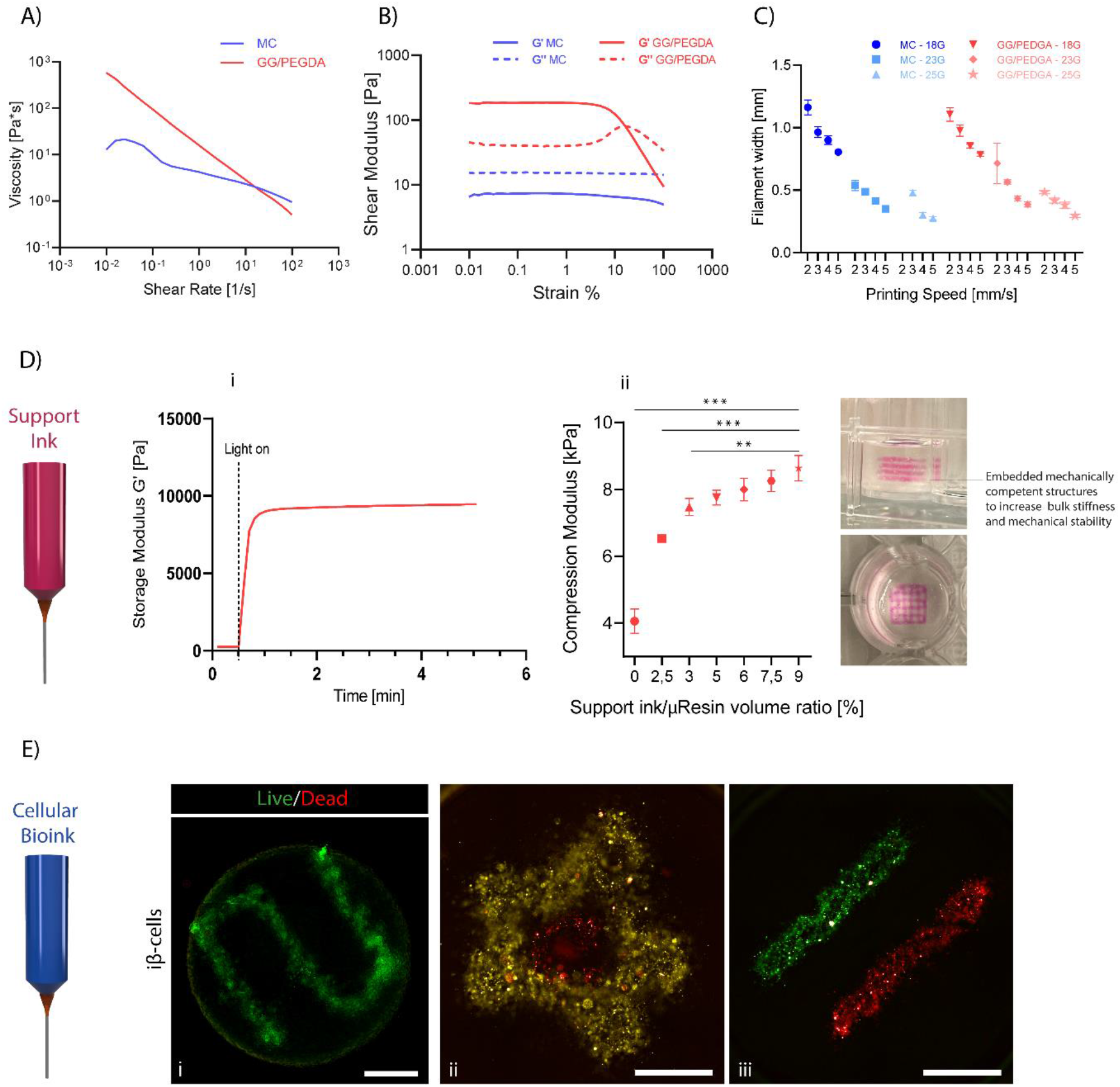
Characterization of the Methylcellulose and GG/PEGDA based bioinks for embedded bioprinting. Representative curves (n=3) for the shear-dependency of ink A) viscosity, and B) storage and loss moduli. C) Printing accuracy of the two formulations, estimated measuring the filament width as a function of the nozzle diameter and printhead translational velocity. D) Support ink: (i) representative (n=3) crosslinking kinetics after light exposure of the GG/PEGDA based bioink and (ii) compressive mechanical properties of the µResin reinforced with GG/PEGDA scaffolds at different volume ratio. E) Cellular bioink: (i) representative live/dead image of extruded iβ-cells at a cell density of 30 million cells/mL (scale bar: 1 mm). (ii,iii) iβ-cells stained with lipophilic membrane dyes, extruded with multiple printheads at a density of 30 million cells/mL (scale bar: 750 µm).

Following the printability optimization, we managed to extrude features of 238 µm and 255 µm with the MC and GG/PEGDA bioinks, respectively (**Figure 5C**), matching the diameter of the smallest needle (250µm) utilized in this study. These values are in the range of the reported resolution of bioinks employed for embedded bioprinting in a granular hydrogel-based support bath.[27] It should be noted that the resolution could be further optimised by the use of different types of support matrices,[68] decreasing the dimension of the microgels and standardizing their shape, while also using smaller extrusion needles.[44,69] We, therefore, tested the combination of GG/PEGDA and the µResin, two materials with significant difference in their storage moduli (**Supplementary Figure S3**), which both could be stabilized via photocrosslinking **(Figure 5Di**).

By patterning and crosslinking different scaffolds with increasing infill within the µResin, we can tune the stiffness of the gels up to increasing the compressive modulus by 108% with a 9% v/v GG/PEGDA concentration (**Figure 5Dii**). After photocrosslinking, the internal GG/PEGDA framework enabled stiffening of the composite structure upon compression, to the point that facile handling with common laboratory spatula was possible without damaging the structural fidelity of the object. Overall, this strategy has relevant implications for the application of embedded printing to create mechanically-competent scaffolds within a granular hydrogel, without compromising the structural properties of the porous granular scaffold, favourable for cell culture.

Finally, we tested whether the extrusion process employing MC was suitable for cell printing. As reported in **Figure 5Ei**, live/dead images of extruded iβ-cells at cell density of 30 ×10^6^ cells/mL showed high viability (⁓100% live cells post printing). By loading multiple cell inks on a multi-printhead bioprinter, it was also possible to generate custom-designed heterocellular compositions, in which each ink is deposited in specific regions (**Figure 5Eii/iii**). Overall, these results on MC and GG/PEGDA inks suggests how embedded printing can be leveraged to produce zonal compartment with defined mechanical and biological composition within granular hydrogels, therefore introducing anisotropy, often necessary in mimicking the hierarchical and specialized architecture of biological tissues. Notably, as the gelMA µResin was demonstrated as independently suitable for spatio-selective annealing via tomographic printing, for the fabrication of highly porous scaffolds and as cell-friendly supporting bath for extrusion bioprinting, the combination of these features constituted the necessary premise for the EmVP process.

### 2.5 Embedded Extrusion Volumetric Printing (EmVP)

To enable the generation of centimetre-scale, geometrically complex constructs laden with precisely patterned multiple cell types, we converged volumetric and extrusion bioprinting into a sequential step process. We showed for the first time the combination of these technologies, which resulted in a hybrid printing technique offering advantages of both the printing approaches, as well as the favourable biological performance of granular hydrogels. According to the recent literature, EmVP could be classified as *Active Null Freeform* 3D printing in the field of freeform three-dimensional printing techniques.[27] “Null” refers to how the support bath is not dissolved and discarded to retrieve the extruded structure, but is kept as part of the printed model, and the “Active” subcategory defines how the extruded bioink and support matrix affect each other.

The EmVP phases are illustrated in **Figure 6**: starting from the design of the models, the features intended to be extruded and the structure to be volumetrically overprinted are saved separately as STL files. Embedded printing was the first printing step: after loading a cylindrical vial with the µResin (either cell-laden or cell-free, depending on the application), the MC bioink was extruded in order to deposit cells in a pre-defined spatial configuration (**Figure 6Ai**). Subsequently, the vial was placed in an in-house developed volumetric printer, where a series of light-projections, delivered at a correct dose, recreated a custom geometry onto the existing part (**Figure 6Aii and iii**). Notably, EmVP further enriches previous composite techniques by sculpting in tens of seconds the microgels in convoluted geometries that cannot be produced by simple molding/casting methods.[52,70] Finally, the vial containing the printed constructs was heated to 37°C to melt the unpolymerized µResin and the sample was washed with prewarmed PBS. The printing process is then followed by the culture and maturation of the living construct.

**Figure 6:**
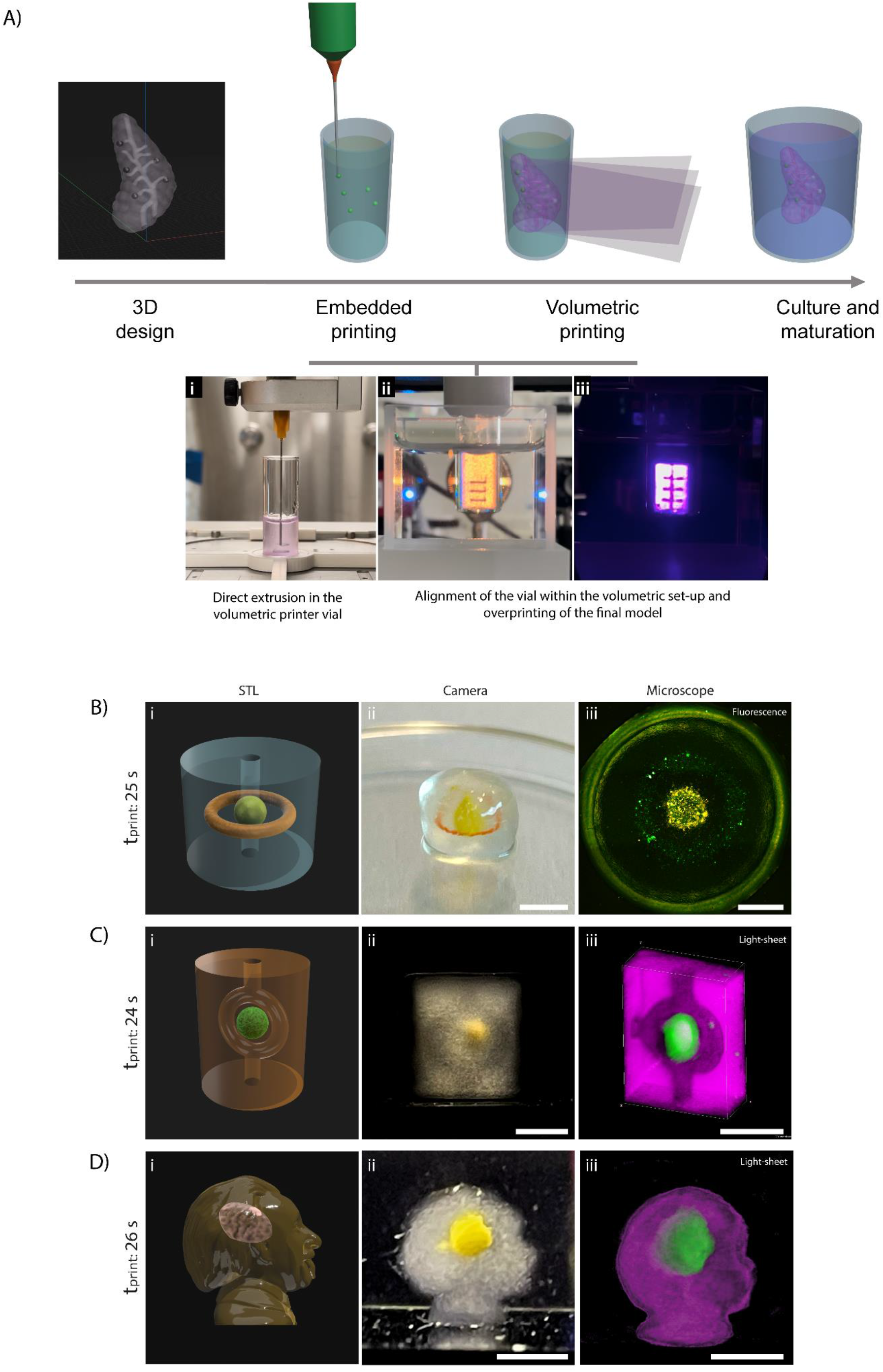
Embedded Extrusion Volumetric Printing (EmVP). A) Sequential steps of the process. (i) The vial containing the µResin is first patterned with the desired cell type and geometry via extrusion bioprinting. Subsequently, (ii) the vial loaded in the volumetric printer (ii) where the light will shape the final structure intertwined with the extruded features (iii). B) Complex co-culture model comprising two different iβ-cells groups (stained with different fluorescent lipophilic membrane dyes), embedded in a volumetrically printed structure with a central channel. Scale bars: ii = 4 mm, iii = 2 mm. C) Example of complex structure model with a branched vasculature; (i) STL design, ii) stereomicrograph, and iii) light-sheet scan reconstructions, digitally cropped to the central region to facilitate visualization of the channels and of the extruded feature. Scale bars: ii = 4 mm, iii = 6 mm. D) Example of complex structure model consisting of the profile of Albert Einstein (with permission of the Albert Einstein Archives, the Hebrew University of Jerusalem. This image cannot be reused in other media or publications), with a central portion of the brain produced by extrusion printing; (i) STL design, ii) stereomicrograph, and iii) light-sheet scan reconstructions. Scale bars: 4 mm.

The potential of EmVP in engineering anisotropic tissues is based on the ability to guide cells fate according to the properties of the microgels, on the precise patterning of multiple cell types, also minimizing the number of cells needed during a volumetric print, with a high-speed over-printing achieved with VBP to potentially use these structures for high-throughput analysis. This new technique addresses a main limitation of the VBP-only approach, which is hampering the broader adoption of this technique: creating more heterogenous constructs, by printing multiple materials and cell types in high densities. This latter condition is indeed difficult to achieve without light-scattering mitigation zstrategies.[5,37] In EmVP, since high-cell densities are produced only in defined, concentrated regions of the vat, we experienced limited printing artefacts, as compared to printing high cell densities (>10^7^ cells/mL), homogenously suspended throughout the bioresins. Hence, in the extrusion printed extruded elements act prevalently as light-attenuating inclusions, rather than scattering particles. While there is likely an upper limit on the volume that can be occupied by highly-cell dense regions, future developments combining computational corrections to counter light attenuation and scattering can be envisioned.

It is important to highlight that embedded, light-absorptive elements have already been shown to be compatible with precise volumetric printing, and more extensive description of the effect of inclusions and light attenuation can be found in other published works.[6,71–73] Given the nature of the tomographic reconstruction algorithm, light profiles can be correctly delivered also in presence of fully opaque elements, as shown with the printing even in presence of a metal rod, as long as an unobstructed 180° view of the inclusion is available.[74] Further underlying this possibility, more recently, volumetric printing in presence of polygonal occlusions with sharp angles was also shown,[73] as well as the correct printing across opaque polycaprolactone-made, melt electrowritten meshes, with light attenuation peaks reported between 10% and 25% of the incoming light intensity.[72] In addition, we experimentally assessed the effect of extrusion printed inclusions at high cell densities, and detected no effect on shape fidelity (**Supplementary Figure S4**).

Printing precision is also dependent on the accuracy of aligning the extruded parts with the projections coming from the DMD. In our set up, the alignment was performed manually. Briefly, light projections were delivered onto the model at a wavelength (green light, 520 nm) far from the excitation spectrum of LAP to avoid premature crosslinking. This enabled the adjustment of the position of the vial to match the starting angle of the tomographic reconstruction. Subsequently, the volumetric printing process was initiated, using a 405nm laser line. Although effective, this method is subject to the expertise of the user, and therefore further automation and standardization are desirable in future iterations, to increase printing resolution and inter-user reproducibility.

Next, we investigated the compatibility of EmVP for the biofabrication of tissue-engineered constructs with complex micro- and macro-geometry, with fast manufacturing times and high degrees of freedom in terms of generating convoluted multicellular shapes. As a proof-of-concept, we first printed dense structures containing a dual culture of extruded iβ-cells in a “Saturn-like” pattern comprising a ring surrounding a printed spheroid, which was finally embedded in a cylindrical volumetric print comprising a central vessel (**Figure 6B**). Similarly, a branched vascular-like channel surrounding a central cluster was also produced by EmVP (**Figure 6C**). Moreover, as proof-of-concept for the production of non-conventional, irregular geometries, a model of Einstein’s head was volumetrically-printed with an inner extrusion printed cluster, shaped and positioned to mimic the midbrain (**Figure 6D**). For all the models, the total volumetric printing time was less than 30 seconds. Additional bioprints comprising of multiple cells types and mimicking a pancreatic model, composed of islet-like cellular structures embedded in a stroma with printed vessels were also produced, as further proof-of-concept (**Supplementary Figure S5**). As the extrusion step is used to pattern only some specific details of the constructs, the total fabrication time, including the alignment procedure remains below 5 minutes. Overall, EmVP still offers advantages over conventional printing strategies in terms of time when printing heterogenous cell-laden centimetre-scale objects in terms of fabrication velocity, (especially when compared to conventional extrusion techniques and DLP processes),[4,75] while also allowing to combine volumetric freedom of design with patterning of high cell density features in a free-form fashion.

### 2.6 EmVP of multicellular synthetic biology-inspired constructs

As a final proof-of-concept experiment, and to assess the functionality of both embedded- and volumetrically-printed cells, we explored the potential of EmVP to create a spatially defined, co-culture system in which a pancreatic cell line interacts with (pre-)adipocytes (**Figure 7**). These cell types were chosen as systems to study their crosstalk are relevant for a variety of metabolic conditions, including diabetes and obesity. Herein, we explored the potential of synthetic biology-based strategies, more specifically using optogenetic tools, to establish engineered and controllable routes of communication between the different (printed) cell types. More specifically, we used iβ-cells, an engineered cell line, which is rewired to store insulin (and the bioluminescent reporter nanoLuc) in intracellular vesicles, and rapidly release its cargo on demand, in response to exposure to visible light (absorption peak at 475 nm), mimicking what native pancreatic β cells do, in response to glucose challenges.[76] This light responsivity is enabled by the induced expression of human melanopsin as photoreceptor, which triggers membrane depolarization and vesicle release upon activation.[76] As adipocyte progenitors, we used hMSCs and explored the effect of the (light-driven) insulin release on their adipogenic differentiation. As shown in Figure 6B, 6C and 6D, complex multi-cellular shapes can be printed with the EmVP process. For this final proof-of-concept experiment, we aimed to provide first evidence that the combined extrusion and volumetric printing process is able to produce constructs capable to express complex cellular functions (i.e. hormone secretion and adipogenic differentiation), which are relevant for future tissue engineering applications. For this, goal, a relatively simple geometry was produced, consisting of a disk shaped construct with an iβ-cells ring at the middle. More precisely, hMSCs were homogenously mixed into the µResin, while iβ-cells were extruded at either 15 or 50×10^6^ cells/mL, to form the circular ring.

**Figure 7:**
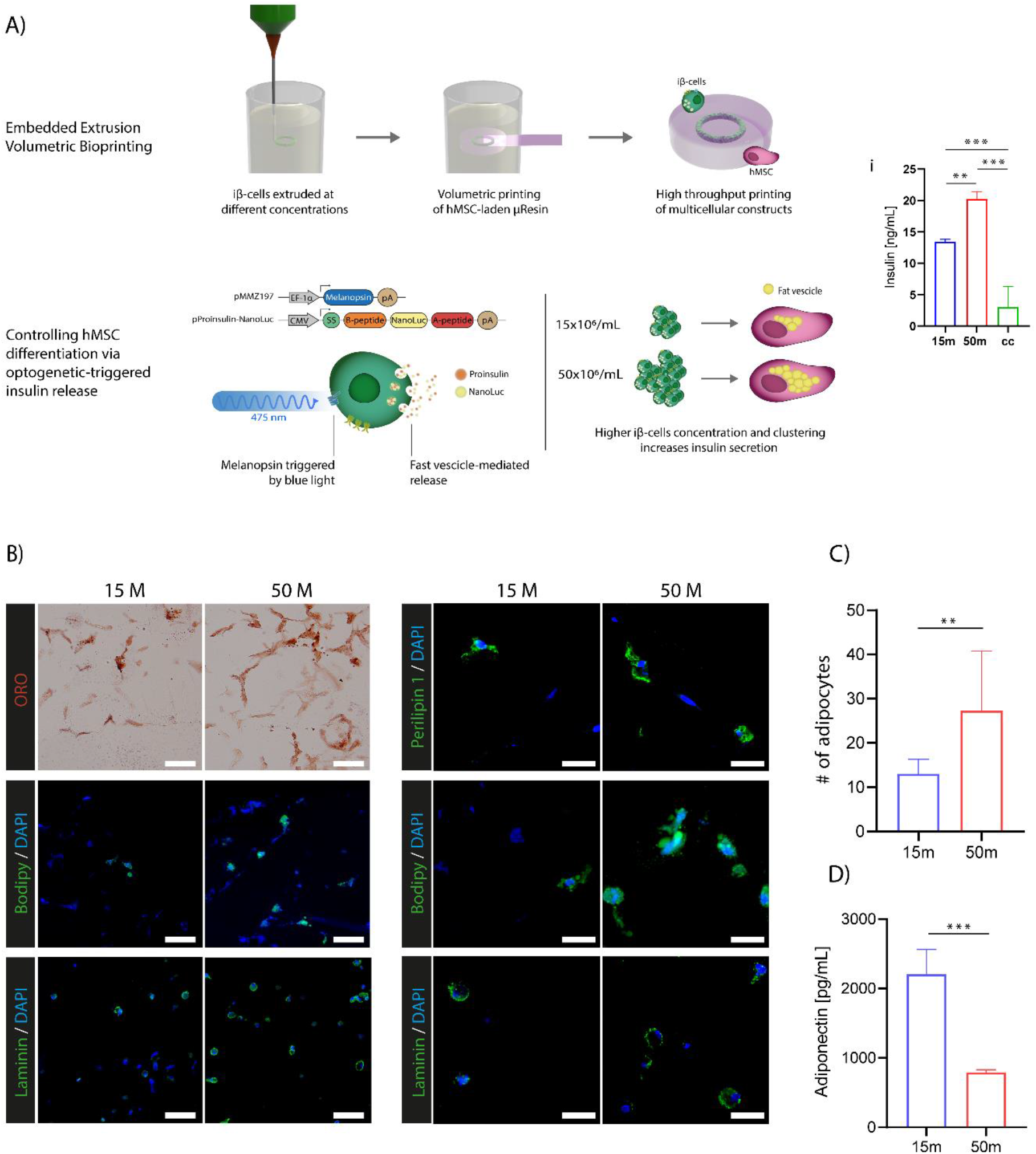
EmVP of multicellular synthetic biology-inspired constructs. A) Schematic representation of the experimental set-up, consisting of extruded rings of iβ-cells at different concentrations embedded in the volumetrically bioprinted gelMA µResin, which is laden with hMSCs. (i) Quantification of the secreted insulin in the supernatant of co-cultured constructs (n=3) over 2 days culture period (timespan between two consecutive culture media refreshments) (cc = 50×10^6^ iβ-cells /mL not extruded). B) Histological staining for fat vesicles using Oil Red O and BODIPY, and immunofluorescent staining for perilipin 1, and laminin, in samples carrying the two EmVP iβ-cells concentrations. Scale bars in B) are: left column = 50 µm, right column = 20 µm. Quantification of the C) amount of differentiated adipocytes (n=6), and of D) adiponectin secreted in the supernatant of samples with the two EmVP iβ-cells concentrations (n=3).

Preliminary steps were taken before the experiments. First, we established that cells undergoing the VBP process maintain a high viability over 7 days of culture (**Supplementary Figure S6**). Then, in another functionality assay, the iβ-cells light-dependent insulin release, already thoroughly characterized in previous works for cells cultured in 2D,[76] was assessed in 3D after bioprinting. A non-zonal co-culture, in which the pancreatic cells (1 million cells/mL, to match the number of iβ-cells per sample to the 50×10^6^/mL extruded condition) were homogenously mixed together with the hMSCs was used as control. Both insulin (Figure 7Aii) and nanoLuc activities (**Supplementary Figure S7**) increased with the iβ-cell content (13.5 ± 0.3 and 20.5 ± 1ng/mL respectively for 15 and 50 million cells/mL). The printed groups outperformed the homogenously distributed co-culture control (3.7 ± 3 ng/mL), suggesting that local high density and possibly cell-cell clustering, enabled by the embedded bioprinting step, are needed for optimal endocrine functionality, in line with what found in previous studies on β-cell like aggregates.[77,78] Having discerned the need to provide close cell-cell contact, we proceeded to assess the effect of the iβ-cells and hMSC interactions after EmVP, over a long-term culture (14 days) in adipogenic media without exogenous insulin. In both groups, hMSCs differentiated adipogenically, as was shown by staining for lipids and perilipin-1. Indeed, perilipin-1 staining was performed as an additional tool to assess the effective differentiation of MSCs into adipocytes, as it is exclusively found on the membrane of lipid droplets [79,80]. The higher iβ-cells concentration (and therefore, higher insulin release), resulted in an increased differentiation of hMSCs into adipocytes. This was qualitatively observed through Oil Red O staining for lipid droplets, as well as from fluorescent staining for lipids using BODIPY, and immunofluorescent staining for the extracellular matrix component laminin, one of the main components of the basal lamina of adipocytes (**Figure 7B**). Moreover, the efficiency of adipogenic commitment was also quantitatively measured from the BODIPY staining, which showed a significantly higher ratio of differentiated adipocytes in the group co-cultured with 50×10^6^ iβ-cells (27 ± 13 % of the cells), compared to the 15×10^6^/mL group (13 ± 3 %) (**Figure 7C**). The adipocyte-specific marker adiponectin, which is secreted by differentiated MSCs, was also detected in the supernatant of both groups. In contrast to the improved adipogenic differentiation found in the group with higher content of iβ-cells, high free-soluble adiponectin was found in the 15×10^6^/mL iβ-cells cultures (2206.9 ± 357 and 790.4 ± 39 pg/mL, for 15×10^6^/mL and 50×10^6^/mL iβ-cells, respectively; **Figure 7D**). In line with this, previous research has also suggested that adiponectin provides a negative feedback for lipid accumulation, and other factors secreted by the iβ-cells could also contribute to this result. Even though a thorough investigation of the determinants of this effect goes beyond the scope of this proof-of-concept assay, and of the objective of this work, which focused on developing the new EmVP biofabrication technology, future experiments can allow to further tune the culture conditions, and probe the mechanistic effect behind pancreatic cell/MSC/adipocyte interactions, with a degree of freedom that is difficult to achieve in *in vivo* models or conventional, non-bioprinted co-cultures. Overall, this pilot study confirms that the EmVPrinted cells (MSC and pancreatic-like line) both preserve their long-term functionality post-printing, opening future possibilities for the generation of complex co-culture model systems, of relevance for a broad array of disease models, besides metabolic dysregulations.

## 3. CONCLUSIONS

Herein, for the first time, the convergence of extrusion-based and volumetric bioprinting provides the possibility to create complex structures on a cm^3^ -scale in a fast and accurate manner, while also introducing multi-material, multi-cellular features in a free-form fashion, a capacity that can greatly expand the range of applications and versatility of volumetric bioprinting. The produced constructs benefit from the mechanically soft, porous microenvironment within the µResin gels, suitable for embedded cells to thrive. Additionally, these soft scaffolds can be easily mechanically reinforced via the in-gel extrusion of structural elements at low-volume fraction. The ability to precisely deposit different cell types at locally high densities and to perform zonal co-cultures, greatly enhances the versatility of volumetrically printed constructs. This enables the biofabrication of composite and “vascularized” structures with spatially varying mechanical and biological characteristics, which is a crucial step in order to better mimic the complex heterogeneous nature of living tissues. Although this study used gelMA as a platform material, the embedded volumetric printing approach can be readily translated to any other photoresponsive, particulate/micro and nano-gel slurry. In future studies, the mixing and even local patterning of microgels obtained from different materials can be envisioned, to create composite scaffolds with unique anisotropic mechanical and degradation properties, or with bioactive pockets releasing drugs or other bioactive compounds within large, bioprinted objects. In addition, the novel concept of combining bioprinting with cells engineered with synthetic biology and optogenetics tools to boost tissue functionality, opens additional opportunities for tissue engineering, regenerative medicine, and the emerging area of engineered living materials.

## 4. EXPERIMENTAL SECTION

### Printable materials

GelMA (85% DoF) was synthesized as previously reported,[81] and dissolved as a 5% w/v solution in phosphate-buffered saline (PBS). Lithium phenyl(2,4,6-trimethylbenzoyl)phosphinate (LAP, Tokyo Chemical Industry, Japan) was added at 0.1% (w/v) as a photoinitiator to induce a photocrosslinking reaction.

Methylcellulose (M0512, Sigma-Aldrich) was dissolved in phosphate-buffered saline (PBS), obtaining a 2 % (w/v) solution to be used as bioink component. A structural biomaterial ink was obtained blending two stock solutions, one obtained dissolving Gellan Gum (GG, Gelzan™ CM, G1910, Sigma-Aldrich) was in deionized water at 80 °C, and the other by dissolving Poly(ethylene glycol) diacrylate (PEGDA, 455008, Sigma-Aldrich) in 10% PBS in deionized water. Both solutions were mixed at 37 °C and LAP was added as a photo-initiator with a final concentration of 0.1 % (w/v). The final GG and PEGDA concentration were 1% w/v and 10% w/v, respectively.

### µResin preparation

The µResin was designed to be used both as support bath for embedded bioprinting and as a photo-crosslinkable material for the volumetric bioprinting process. The process to produce milled GelMA microgels, was adapted from a previously published protocol.[11] Sterile, lyophilized porcine GelMA was weighed and dissolved in PBS at a concentration of 5% (w/v) and subsequently gelated at 4 °C overnight. A 500 ml Oster® blender mason jar, UV sterilized and also stored at 4 °C overnight, was filled with cold PBS to make up a total volume of 100 ml with the solid block of gelated GelMA. The GelMA was milled for different blending times (see Figure 2) at “pulse” speed and the contents of the jar were centrifuged at 4000 rpm for 4 mins at 4°C to separate the GelMA microparticles from the PBS. After centrifugation, the supernatant was aspirated and, in case bubbles were present, the milled GelMA was washed with cold PBS and centrifuged again until all bubbles were removed. With the repeated centrifugation, bubble and supernatant removal steps, the GelMA particles are pelleted and the same process is performed across different batches to ensure consistency in the packing density. The GelMA slurry was stored at 4 °C until further use.

### Mechanical testing

Compressive properties of the µResins gels were assessed via a uniaxial, unconfined compression test, with a Dynamic Mechanical Analyzer (DMA Q800, TA Instruments, The Netherlands), equipped with a cylindrical flat piston. Cylindrical samples (diameter = 6 mm, height= 2 mm) were prepared by casting the µResin slurry in a mold and crosslinked upon exposure to a 365 nm light source (5 minutes). Samples (n=4) were left overnight at 37°C, and subsequently subjected to a force ramp 0.5 N min^−1^. The compression modulus was calculated as the slope of the stress/strain curve in the 10–15% strain range.

### Rheological evaluation

All rheological measurements were performed using a Discovery HR-2, TA Instrument rheometer. Shear rate tests were performed on µResins and on both the MC and GG/PEGDA bioinks (n=3) using a 40mm parallel plate geometry (0.2 mm gap). Viscosity and shear stress were recorded during a shear rate ramp between 0 and 100 s-1. Strain sweep was performed on both bioinks (n=3) using a 40mm parallel plate geometry (0.2 mm gap). The storage (G′) and loss (G″) moduli were recorded during a strain % ramp between 0.01 and 100%. Photorheological analysis was performed on µResins and on the GG/PEGDA ink (n=3) using a 20mm parallel plate geometry (0.3 mm gap), by outfitting the rheometer with a UV light source (Vilber Lourmat UV Lamp, Collegien, France). Photopolymerization was initiated through exposure of the gels to light after 30 seconds, and light was kept on for 3 minutes. The storage (G′) and loss (G″) moduli were recorded during a time sweep of 5 minutes.

### Porosity and microparticles characterization

Porosity characterization was performed on annealed2 gels stained with Cyanine-3.5 dye (Cy3.5). Samples were washed with PBS and imaged through a custom-built light sheet microscope. Porosity was determined on single cross-section images and using a threshold to select the void spaces via ImageJ. Microgels size analysis was performed by dispersing microgels on a glass slide and imaging with an optic microscope (Olympus SZ61). For each condition, the diameter of microgels (n≥300) was measured and analyzed using ImageJ.

### Scattering quantification

Light scattering effect as a function of the temperature was characterized with a custom-made optical setup. A laser beam (532 nm, Roithner RLDD532-50-3) having a circular cross-section was shined through the centre of a PMMA squared (10 mm x 10 mm) cuvette containing the µResin, and the resulting scattering pattern was projected on a camera photodetector (Olympus E-PL7, lens removed) placed 15 cm behind the cuvette. The full width half maximum (FWHM) of the patterns was then calculated from the center along the horizontal axes with a MATLAB script on different pictures taken at different time points (every 20 seconds) as the µResin samples (n=3) slowly warmed up to room temperature. A thermal camera (Testo 880) was placed on top of the setup allowed to monitor the temperature of the µResin during the measurements

### Cells isolation and culture

GFP-labeled human umbilical vein endothelial cells (GFP-HUVECs; cAP-001GFP, Angioproteomie) were expanded in Endothelial Cell Growth Medium-2 (EGM-2, Lonza), containing Endothelial Basal Medium-2 + SingleQuots (Lonza), supplemented with 10% fetal bovine serum (FBS) and 1% penicillin/streptomycin (P/S). Culture flasks were pre-coated with 0.1% Gelatin for 30 minutes at 37°C, followed by 3 washes with phosphate-buffered saline (PBS). GFP-HUVECS were used in experiments at passage 6-7.

Human bone marrow-derived mesenchymal stromal cells (MSCs) were isolated from bone marrow aspirates of consenting patients, as previously described.[82] Briefly, human bone marrow aspirates were obtained from the iliac crest of patients that were receiving spondylodesis or hip replacement surgery. Isolation and distribution were performed in accordance with protocols approved by the Biobank Research Ethics Committee (isolation 08-001, distribution protocol 18-739, University Medical Center Utrecht). Protocols used are in line with the principles embodied in the Declaration of Helsinki. MSCs were expanded in ⍺-Modified Eagle Medium culture medium (α-MEM, Gibco™, Life Technologies) supplemented with 10% FBS, 1% P/S, 1% L-ascorbic acid-2-phosphate (ASAP; Sigma-Aldrich, The Netherlands) and 1 ng/ml basic fibroblast growth factor (bFGF; R&D Systems) and used at passage 4-5.

For adipogenic differentiation of the MSCs, medium consisted of ⍺-MEM (Gibco™, Life technologies) supplemented with 1% P/S, 0.4 µg/mL dexamethasone (Sigma-Aldrich, The Netherlands), 0.052 mM 3-isobutyl-1-methylxanthine (IBMX; Sigma-Aldrich, The Netherlands), 0.2 mM indomethacin (Sigma-Aldrich, The Netherlands). When indicated, adipogenic medium was supplemented with 10 µg/mL insulin (Sigma-Aldrich, The Netherlands).

LUHMES were expanded and differentiated as previously described.[83] Briefly, neural differentiation medium was used, which contains the antibiotic doxycycline (dox) to inhibit gene expression via the tet-off system, a genetic manipulation technique that is used to spatially and temporally control gene expression.[83] Other supplements included dibutyryl-cyclic-AMP (db-cAMP) to stimulate neurite outgrowth, neural differentiation and maturation[83–85] and glial cell- and brain-derived neurotrophic factors (GDNF and BDNF) to promote survival of and induce differentiation to dopaminergic neurons. After 2 days of differentiation in 2D monolayer conditions, LUHMES cells were harvested from culture flasks by detaching with 3 mL of 0.25% Trypsin+EDTA for 7 min at 37°C.

iβ-cells were obtained and cultured as previously described.[76,86] Briefly, a 1.1E7 cell line-derived cell clone deficient in glucose-sensitive insulin secretion was transduced with Proinsulin-NanoLuc-derived lentiviral particles and selected in culture medium containing 5 µg/mL blasticidin to create polyclonal INSvesc cells. Next, a monoclonal cell population, was picked based on the best performance for depolarization-triggered nanoLuc secretion. This cell line was co-transfected with the SB100X expression vector pCMV-T7-SB100 (PhCMV-SB100X-pA) and the SB100X-specific transposon pMMZ197 (ITR-PhEF1α-melanopsin-pA-PRPBSA-ypet-P2A-PuroR-pA-ITR) to generate a polyclonal population of iβ-cells that stably expressed melanopsin as well as Proinsulin-NanoLuc cassettes. After selection for two weeks in medium containing 5 µg/mL blasticidin and 1 µg/mL puromycin, the monoclonal iβ-cells were sorted by means of FACS (Becton Dickinson LSRII Fortessa flow cytometer) and screened for blue-light-responsive nanoLuc secretion. iβ-cell were cultured in RPMI 1640 Medium, GlutaMAX™, HEPES (Gibco™, Life technologies) supplemented with FBS (10% v/v) and 1% P/S and used at passage 3-4.

All cells were cultured in a 95% humidified incubator at 37°C, 5% CO2.

### Volumetric printing

A custom-built volumetric printer was used for the volumetric printing step. The optical system was comprised of a 405nm 1W diode laser (Ushio HL40033G) mounted on a temperature-controlled mount (Thorlabs LDM90/M), and a digital micromirror device (DMD) (Vialux v7000 HiSpeed). A cylindrical vial (to be used as the cylindrical print volume) of inner diameter 13.2mm was filled with the µResin, and vertically suspended within the rotary stage. The vial was also immersed within a water-filled 40mm square cuvette (Hellma 704.002-OG) containing an index-matching liquid needed to minimise distortions resulting from the curvature of the vial walls. A MATLAB code was used to operate the device. After printing, the thermally gelled µResin was gently washed with pre-warmed PBS at 37° C to remove the uncrosslinked material from the printed structures.

### Extrusion (bio)printing

Three-dimensional (bio)printing experiments were performed with a 3D Discovery bioprinter (regenHu, Switzerland), using a pneumatic-driven extrusion printhead. For printing evaluation, 3 mL syringes were loaded with 1 mL of Cy-3.5 stained bioink (both for GG/PEGDA and MC), three different needles (18G, 23G and 25G) and three different printing speeds (2, 3, 4 and 5 mm/s) were tested to characterize the filaments width. Images (n = 5) of S-shape embedded filaments were taken with a confocal microscope (SPX8, Leica Microsystems, The Netherlands) and analyzed with ImageJ. For mechanical reinforcement of µResin gels, GG/PGEDA based filaments were printed at 5 mm/s speed with a 25G needle (250 µm diameter) in different S-shape layers.

For cell viability after printing, iβ-cells were printed at 5 mm/s speed with a 25G needle (250 µm diameter), and cell viability was evaluated using a live/dead assay (calcein AM/ethidium homodimer, Thermo Fischer Scientific, The Netherlands) on gels (n = 3).

For multicellular extrusion printing (Figure 5Cii/iii) iβ-cells were pre-stained (Vybrant™ Multicolor Cell-Labeling Kit, Thermo Fischer Scientific, The Netherlands) printed at 5 mm/s speed with a 25G needle (250 µm diameter). All G-codes were manually written.

### Embedded Volumetric Printing (EmVP)

The features meant to be extruded and the structure to be volumetrically overprint were designed and saved separately as STL files. First, a cylindrical vial was loaded with the µResin at 4°C (either cell-laden or cell-free, depending on the application) + 0.1% w/v LAP, everything was then gently mixed directly in the printing vial. The vial was then placed in a 3D Discovery extrusion printer (regenHu, Switzerland) and kept in place through a custom printed holder. 3 mL syringes were loaded with 1 mL of (bio)ink and the G-codes of the extruded features were manually written. Subsequently, keeping the vial protected from light exposure to prevent the crosslinking of the µResin, the vial was placed into a custom-built volumetric printer. To align the extruded features with the model intended to be overprinted, a first series of light-projections was delivered on the rotating vial with a 520 nm laser, in order to prevent the crosslinking during this calibration process. Therefore, light-projections of the model meant to be volumetrically printed were delivered with a 405 nm laser, until the curing of the µResin was achieved. Finally, the vial was heated to 37°C to melt the unpolymerized µResin and the sample was washed with prewarmed PBS.

### LipidTox, Phalloidin and βIII-tubulin staining

Gels were submerged in 0.1% Triton-100 and incubated at room temperature for 15 mins to allow permeabilization of the cell membrane. To prevent non-specific binding of antibodies to membrane proteins, gels were washed 3 times x 5 minutes with a 3% Bovine Serum Albumin (BSA, Sigma) blocking solution in PBS. To stain βIII-tubulin on LUHMES laden gels, a primary mouse antibody (1:500) was used and gels were stained in the dark at 4°C overnight. After the incubation period, the samples were washed at low rocker speed with PBS 2 times for 5 hrs. Next, secondary antibodies diluted in the BSA solution were added to the samples and incubated in the dark at 4°C overnight, using goat antimouse IgG AlexaFluor 488 (Invitrogen; 2mg/ml). Following this step, the nuclei were stained at room temperature for 15 minuts with 4’,6-diamidino-2-phenylindole (DAPI)(Sigma Aldrich, D9542, 1:500, 1 mg/ml) to yield a blue nucleic fluorescence by binding to nucleic acids.

To stain phalloidin on MSCs laden gels, phalloidin (Invitrogen, Alexa Fluor 488 Phalloidin) (1:200) was used, and gels were stained in the dark at room temperature for 30 minutes. Nuclei were stained at room temperature for 3 minutes with DAPI (Sigma Aldrich, D9542, 1:500, 1 mg/ml).

To stain lipid vesicles on differentiated adipocytes, LipidTox (Invitrogen, HCS LipidTOX Green Neutral Lipid Stain)(1:200) was used, and gels were stained in the dark at room temperature for 3 hours. Nuclei were stained at room temperature for 3 minutes with DAPI (Sigma Aldrich, D9542, 1:500, 1 mg/ml). All samples were washed with PBS and covered with aluminium foil until imaging.

### Oil red O and BODIPY staining

Gels were embedded in Tissue-Tek (Sakura Finetek, Torrance, CA, USA) and incubated overnight in a wet chamber at room temperature. The samples were snap-frozen in liquid nitrogen the following day, cut into 6 μm sections using a cryostat (CM 3050S, Leica, Wetzlar, Germany), and transferred onto microscope slides. The slides were then soaked in ddH_2_O for 30 s to dissolve residual Tissue-Tek®. To visualize lipid content, sections were stained with Oil Red O (3 mg/mL Oil Red O in 60% isopropanol; Sigma-Aldrich, St. Louis, MO, USA) for 6 minutes. After staining, the slides were washed under cold running water for 10 min and then coverslips were mounted with glycergel (Dako, Hamburg, Germany). As an additional lipid visualization, sections were stained with BODIPY 493/503 (ThermoFisher, Waltham, MA, USA) diluted 1:500 in PBS for 30 min at room temperature, washed three times with PBS and mounted with DAPI mounting medium ImmunoSelect (Dako, Hamburg, Germany). Images were taken with a fluorescence microscope (Olympus BX51/DP71).

### Perilipin 1 and Laminin staining

Antigen retrieval was performed using proteinase K digestion for 10 min at room temperature, followed by three washing steps with PBS + 0.2% Triton-×100. For perilipin staining, samples were incubated in citric acid buffer (10X citric acid buffer pH 6.0, ab64214, Abcam, Cambridge, UK) at 100°C for 5 min, cooled at RT for 20 minutes and washed with PBS. Cryosections were blocked with 1% bovine serum albumin in PBS for 1 h at room temperature and incubated with primary antibodies (anti-perilipin 1, PA5-72921, 1:200, ThermoFisher, Waltham, MA, USA; anti-laminin, ab11575, 1:200, Abcam, Cambridge, UK) diluted in 1% BSA in PBS. After incubation in a humidified chamber overnight at room temperature, cryosections were washed three times with PBS and incubated with the secondary antibody (goat anti-rabbit Alexa488, 111-545-144, 1:400, Biozol, Eching, Germany; diluted in 1% BSA in PBS) in a darkened humidified chamber for 1 h at room temperature. After three additional washing steps in PBS, sections were mounted with DAPI mounting medium ImmunoSelect (Dako, Hamburg, Germany), and images were taken with a fluorescence microscope (Olympus BX51/DP71 and Zeiss Colibri7) and post-processed with Olympus CellSense Dimensions Software.

### Adiponectin and insulin ELISA

Adiponectin concentrations were determined using human adiponectin/Acrp30 DuoSet ELISA (R&D Systems, Minneapolis, MN, USA) following the manufacturer’s instructions. The levels of mouse insulin in culture supernatants were quantified with a Rat/Mouse Insulin ELISA kit (Mercodia; cat. no.10-1232-01) following the manufacturer’s instructions.

### Statistical Analyzes

Results were reported as mean ± standard deviation (S.D.). Statistical analysis was performed using GraphPad Prism 8.0.2 (GraphPad Software, USA). Comparisons between multiple (> 2) experimental groups were assessed via one or two-way ANOVAs, followed by post hoc Bonferroni correction to test differences between groups. Student’s t-test was performed between 2 experimental groups for statistical analysis. Unpaired t-test was used for parametric comparisons of data sets. Non-parametric tests were used when normality could not be assumed. Differences were considered significant when p < 0.05. Significance is expressed on graphs as follows: * p <= 0.05, ** p <= 0.01, *** p <= 0.001, **** p <= 0.0001.

## AKNOWLEDGMENTS

This project received funding from the European Research Council (ERC) under the European Union’s Horizon 2020 research and innovation programme (grant agreement No. 949806, VOLUME-BIO) and from the European’s Union’s Horizon 2020 research and innovation programme under grant agreement No 964497 (ENLIGHT). R.L and J.M acknowledge the funding from the ReumaNederland (LLP-12, LLP22, and 19-1-207 MINIJOINT) and the Gravitation Program “Materials Driven Regeneration”, funded by the Netherlands Organization for Scientific Research (024.003.013). R.L. also acknowledges funding from the NWA-Ideeëngenerator programme of the Netherlands Organization for Scientific Research (NWA.1228.192.105). T.B. acknowledges the funding from the Deutsche Forschungsgemeinschaft (DFG, German Research Foundation), Project number 326998133, TRR 225 (subproject C02).

## Supporting information

### SUPPLEMENTARY FIGURES

**Supplementary Figure S1:**
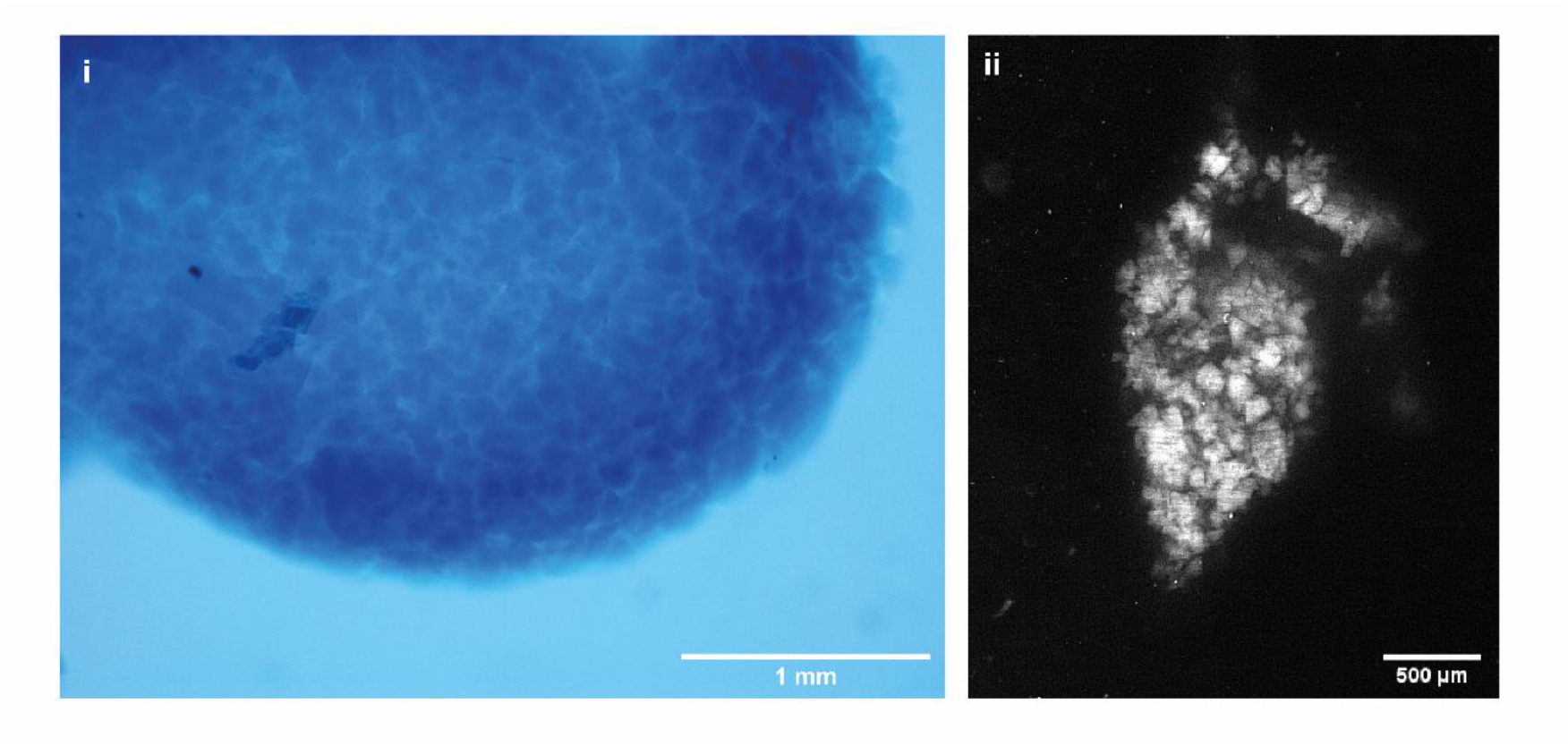
Granular constructs characterised by irregular microparticles. i) Optical microscopy image of a volumetric printed constructs, stained with Trypan blue to facilitate visualization, showing the particulate composition. ii) Light-sheet imaging of the irregularly-shaped microparticles.

**Supplementary Figure S2:**
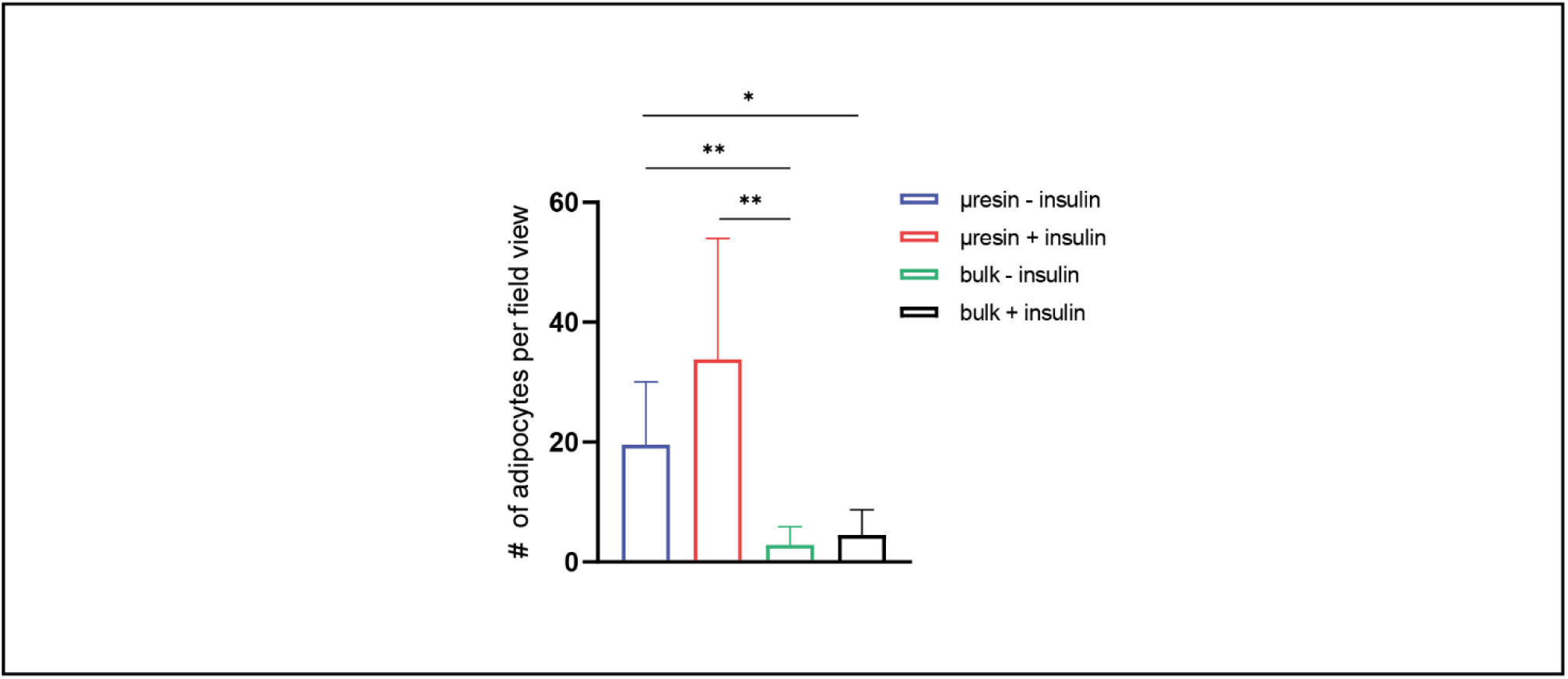
Effect of the microstructure of the microgel based gels on hMSC cells differentiation into adipocytes. Quantification of the number of adipocytes per field view in bulk and gelMA µResin, cultured with and without insulin in the differentiation medium.

**Supplementary Figure S3:**
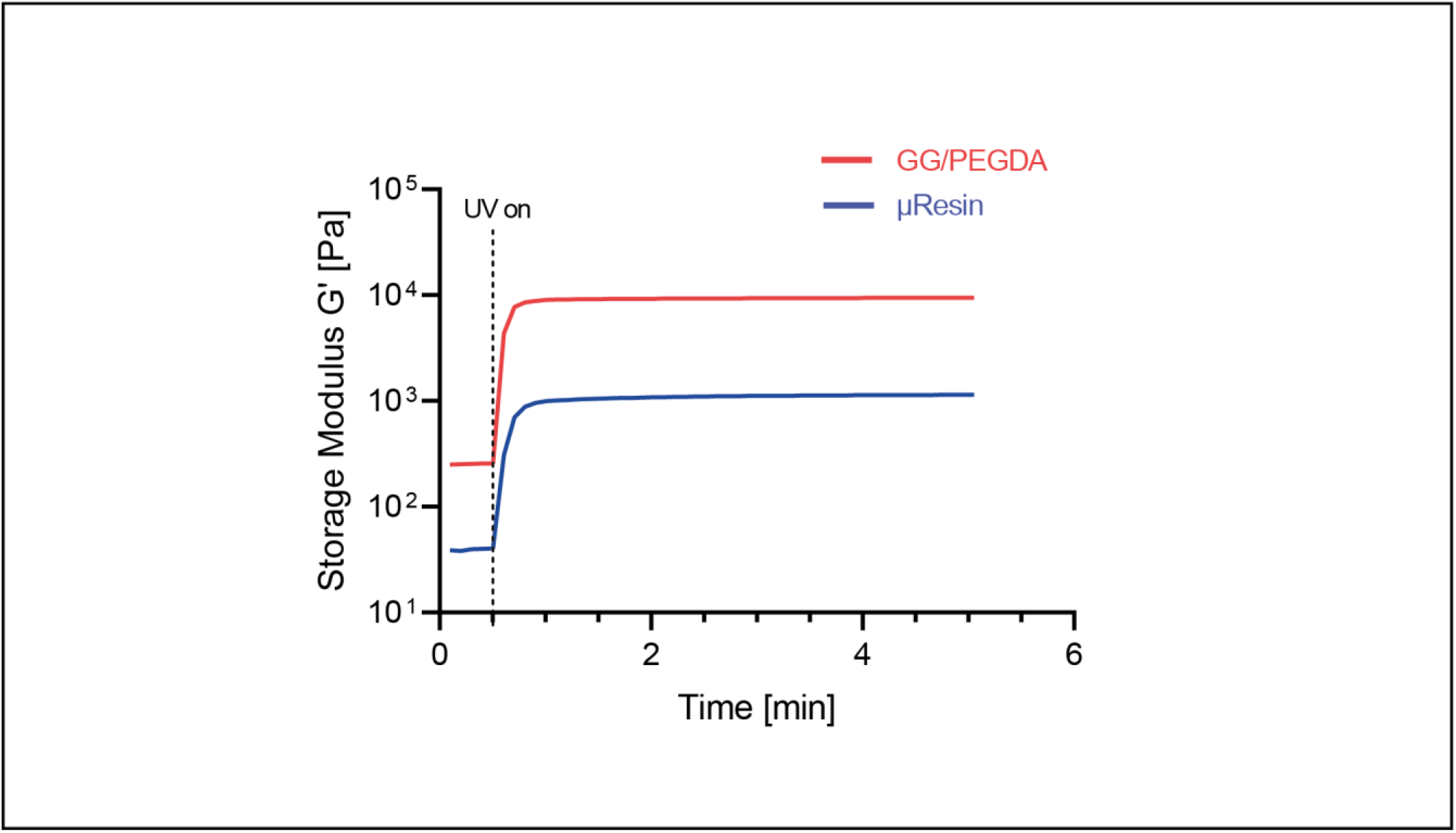
Representative (n=3) photorheologies of GG/PEGDA ink and µResin.

**Supplemetary Figure S4:**
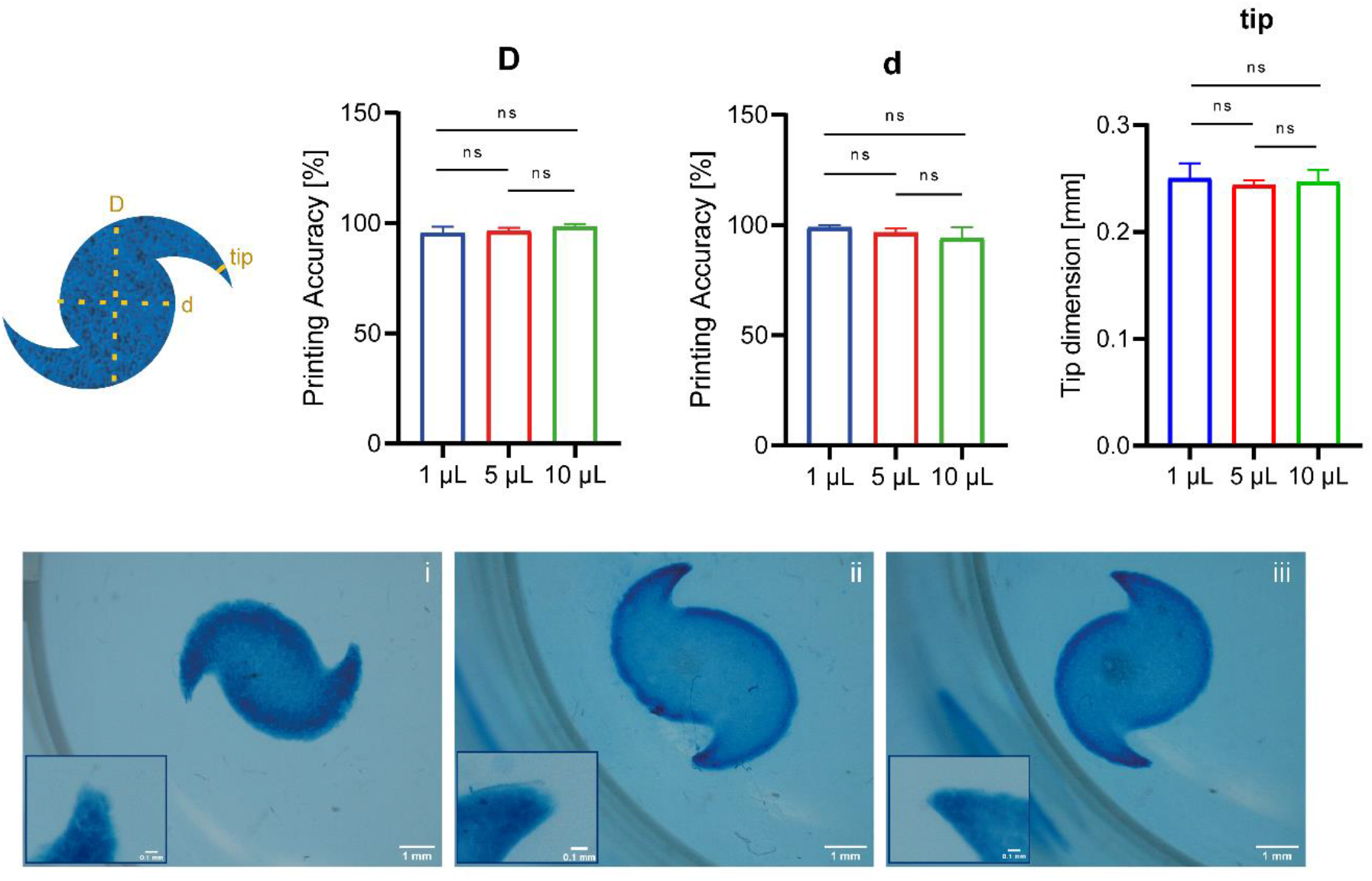
Effect of the extruded features on the volumetric printing process and on the resolution of the final construct. Printing accuracy (n=5), calculated on the bigger (D) and smaller (d) diameters of the galaxy-shape model, and dimensions comparison of the galaxy-tips, as a function of the different volumes of iβ-cells-laded bioink extruded in the middle: i = 1 µL; ii = 5 µL; iii = 10 µL. iβ-cells concentration: 30×10^6^ cells/mL. Scale bars: overviews = 1 mm; magnifications = 0.1 mm.

**Supplementary Figure S5:**
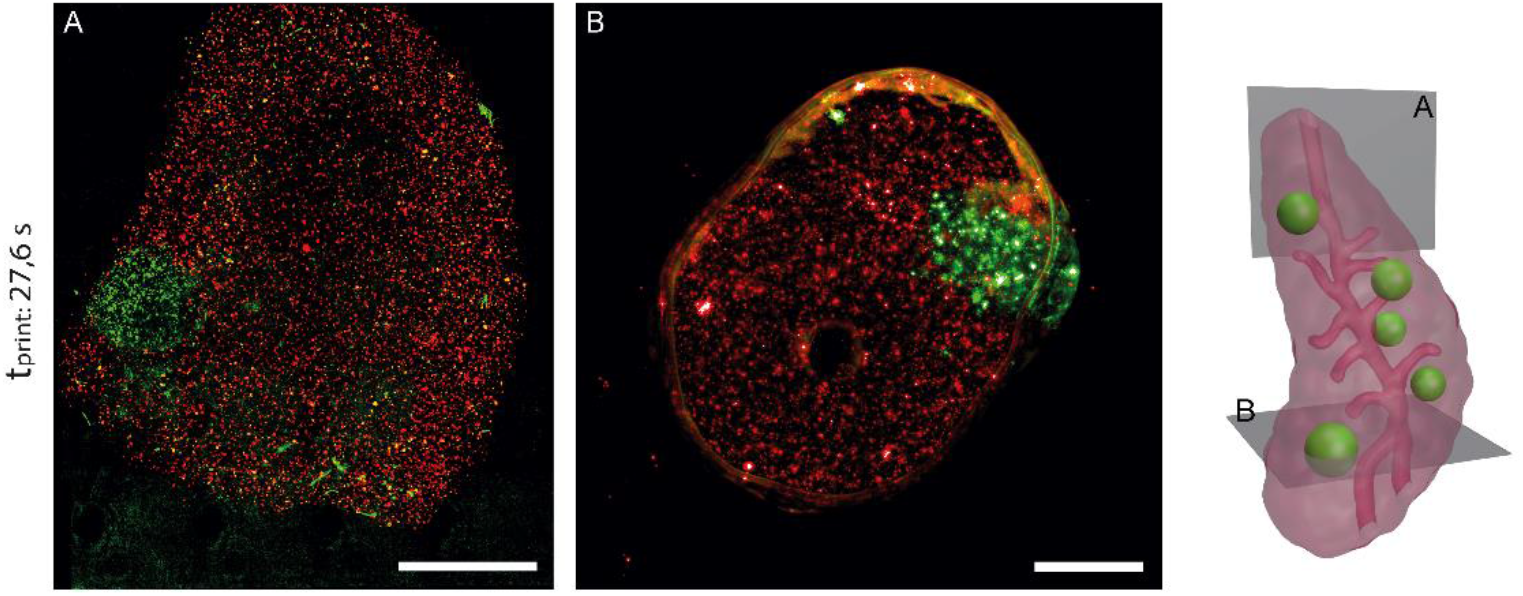
Complex multicellular 3D construct representing a pancreas model were produced by mixing the red-labelled stromal compartment (MSC, Vybrant DiD membrane staining), and via extrusion printing of clusters of green-labelled pancreatic cells (iβ-cells, Vybrant DiO). Tomographic light projections were applied to volumetrically print the outer pancreas shape and the inner channels representing the main branches of the vasculature. A) and B) show two randomly selected confocal scanning fluorescence microscopy cross sections with an open printed channel visible in B). Scale bars: A = 500 µm; B = 1 mm.

**Supplementary Figure S6:**
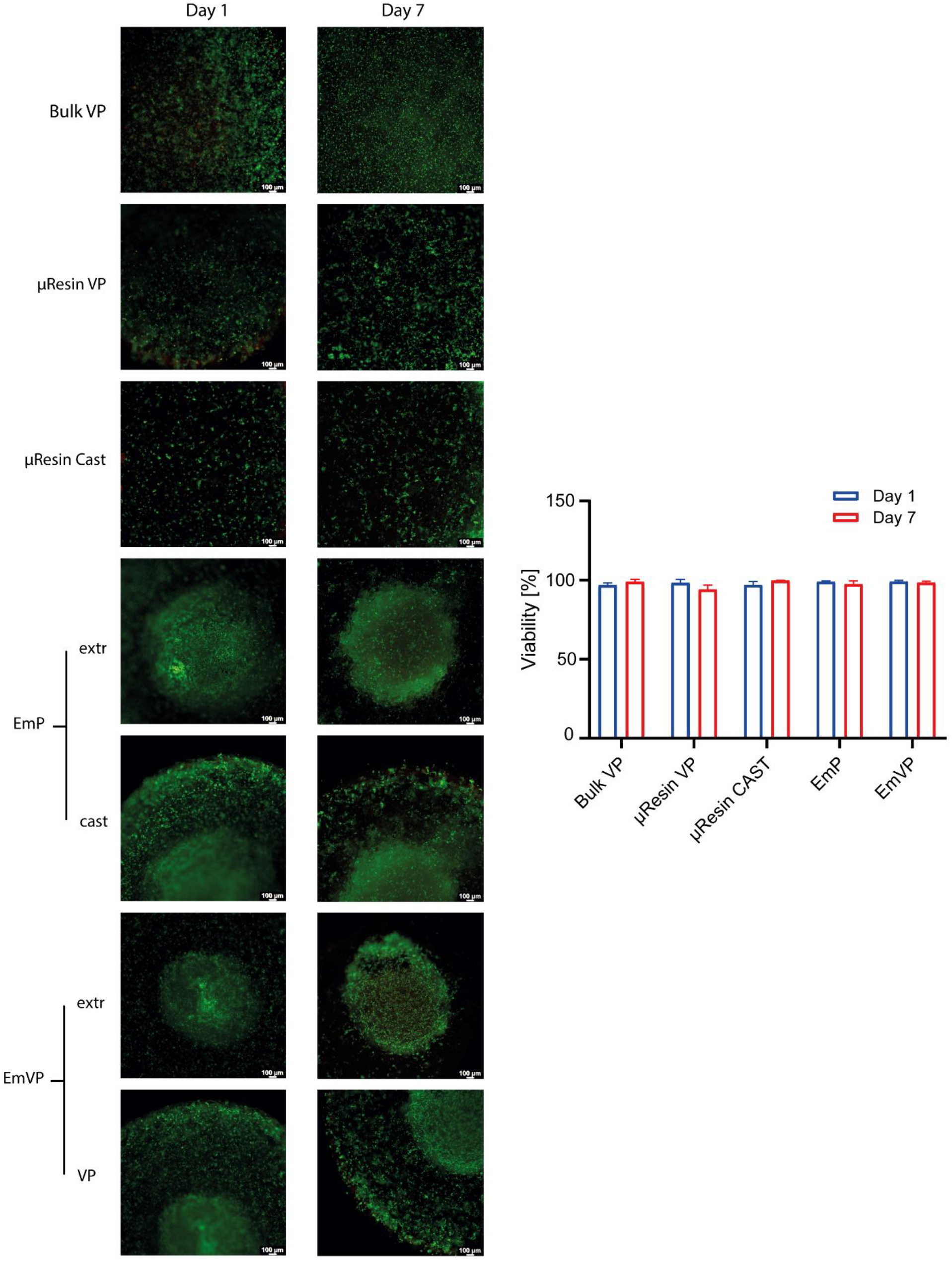
Viability quantification on Live/Dead images at day 1 and day 7 (n=3) of iβ-cells bioprinted with different approaches: volumetric bioprinted in bulk gelMA (Bulk VP) at a cell density of 2.5 million cells/mL. Volumetric bioprinted in µResin (µResin VP) at a cell density of 2.5 million cells/mL. Casted µResin (µResin Cast) at a cell density of 2.5 million cells/mL. Extruded at a cell density of 30 million cells/mL (EmP – extr) in a 2.5 million cells/mL-laden and casted µResin (EmP – cast). Extruded at a cell density of 30 million cells/mL (EmVP – extr) in a 2.5 million cells/mL-laden and volumetric printed µResin (EmVP – VP). The latter condition represent the EmVP process described in the manuscript.

**Supplementary Figure S7:**
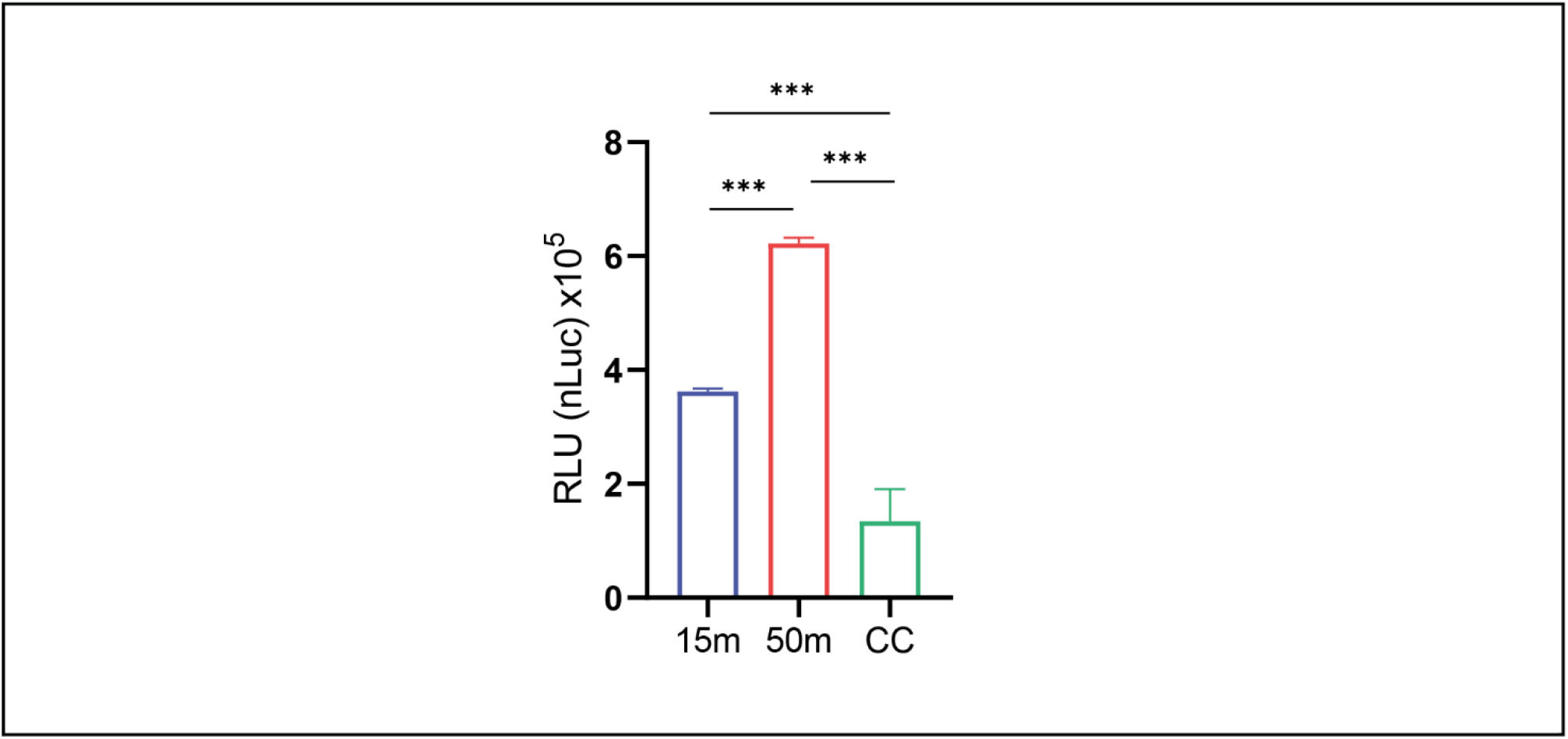
NanoLuc levels in cell culture supernatants (n=3) of the extruded and mixed iβ-cells quantified using the Nano-Glo Luciferase Assay System (N1110, Promega) (cc = 50×10^6^ iβ-cells /mL not extruded).

## REFERENCES

[1] L. Moroni, T. Boland, J. A. Burdick, C. De Maria, B. Derby, G. Forgacs, J. Groll, Q. Li, J. Malda, V. A. Mironov, C. Mota, M. Nakamura, W. Shu, S. Takeuchi, T. B. F. Woodfield, T. Xu, J. J. Yoo, G. Vozzi, Trends in Biotechnology 2018, 36, 384.

[2] H.-W. Kang, S. J. Lee, I. K. Ko, C. Kengla, J. J. Yoo, A. Atala, Nat Biotechnol 2016, 34, 312.

[3] R. Levato, T. Jungst, R. G. Scheuring, T. Blunk, J. Groll, J. Malda, Advanced Materials 2020, 32, 1906423.

[4] P. N. Bernal, P. Delrot, D. Loterie, Y. Li, J. Malda, C. Moser, R. Levato, Adv. Mater. 2019, 31, 1904209.

[5] P. N. Bernal, M. Bouwmeester, J. Madrid-Wolff, M. Falandt, S. Florczak, N. G. Rodriguez, Y. Li, G. Größbacher, R. Samsom, M. van Wolferen, L. J. W. van der Laan, P. Delrot, D. Loterie, J. Malda, C. Moser, B. Spee, R. Levato, Advanced Materials 2022, 34, 2110054.

[6] B. E. Kelly, I. Bhattacharya, H. Heidari, M. Shusteff, C. M. Spadaccini, H. K. Taylor, Science 2019, 363, 1075.

7. D. Loterie, P. Delrot, C. Moser, Volumetric 3D Printing of Elastomers by Tomographic Back-Projections, 2018.

[8] J. Groll, J. A. Burdick, D.-W. Cho, B. Derby, M. Gelinsky, S. C. Heilshorn, T. Jüngst, J. Malda, V. A. Mironov, K. Nakayama, A. Ovsianikov, W. Sun, S. Takeuchi, J. J. Yoo, T. B. F. Woodfield, Biofabrication 2018, 11, 013001.

[9] K. Flégeau, A. Puiggali-Jou, M. Zenobi-Wong, Biofabrication 2022, 14, 034105.

[10] A. Sinclair, M. B. O’Kelly, T. Bai, H.-C. Hung, P. Jain, S. Jiang, Advanced Materials 2018, 30, 1803087.

[11] T. J. Hinton, Q. Jallerat, R. N. Palchesko, J. H. Park, M. S. Grodzicki, H.-J. Shue, M. H. Ramadan, A. R. Hudson, A. W. Feinberg, Science Advances 2015, 1, e1500758.

[12] D. M. Headen, J. R. García, A. J. García, Microsyst Nanoeng 2018, 4, 1.

[13] L. Zhang, K. Chen, H. Zhang, B. Pang, C.-H. Choi, A. S. Mao, H. Liao, S. Utech, D. J. Mooney, H. Wang, D. A. Weitz, Small 2018, 14, 1702955.

[14] G. C. Le Goff, J. Lee, A. Gupta, W. A. Hill, P. S. Doyle, Advanced Science 2015, 2, 1500149.

[15] W. Yang, H. Yu, G. Li, Y. Wang, L. Liu, Small 2017, 13, 1602769.

[16] W. Leong, T. T. Lau, D.-A. Wang, Acta Biomater 2013, 9, 6459.

[17] C. L. Franco, J. Price, J. L. West, Acta Biomaterialia 2011, 7, 3267.

[18] J. E. Mealy, J. J. Chung, H.-H. Jeong, D. Issadore, D. Lee, P. Atluri, J. A. Burdick, Advanced Materials 2018, 30, 1705912.

[19] T. H. Qazi, J. A. Burdick, Biomaterials and Biosystems 2021, 1, 100008.

[20] A. R. Anderson, E. Nicklow, T. Segura, Acta Biomaterialia 2022, 150, 111.

[21] E. Sideris, S. Kioulaphides, K. L. Wilson, A. Yu, J. Chen, S. T. Carmichael, T. Segura, Advanced Therapeutics 2022, 5, 2200048.

[22] A. C. Daly, L. Riley, T. Segura, J. A. Burdick, Nat Rev Mater 2020, 5, 20.

[23] L. Riley, L. Schirmer, T. Segura, Curr Opin Biotechnol 2019, 60, 1.

[24] R.-S. Hsu, P.-Y. Chen, J.-H. Fang, Y.-Y. Chen, C.-W. Chang, Y.-J. Lu, S.-H. Hu, Advanced Science 2019, 6, 1900520.

[25] S. Xin, D. Chimene, J. E. Garza, A. K. Gaharwar, D. L. Alge, Biomater. Sci. 2019, 7, 1179.

[26] C. B. Highley, K. H. Song, A. C. Daly, J. A. Burdick, Advanced Science 2019, 6, 1801076.

[27] A. Colly, C. Marquette, J.-M. Frances, E.-J. Courtial, MRS Bulletin 2022, DOI 10.1557/s43577-022-00348-9.

[28] D. J. Shiwarski, A. R. Hudson, J. W. Tashman, A. W. Feinberg, APL Bioengineering 2021, 5, 010904.

[29] K. Zhou, Y. Sun, J. Yang, H. Mao, Z. Gu, J. Mater. Chem. B 2022, 10, 1897.

[30] A. Schwab, R. Levato, M. D’Este, S. Piluso, D. Eglin, J. Malda, Chem. Rev. 2020, 120, 11028.

[31] T. Bhattacharjee, S. M. Zehnder, K. G. Rowe, S. Jain, R. M. Nixon, W. G. Sawyer, T. E. Angelini, Science Advances 2015, 1, e1500655.

[32] S. Raees, F. Ullah, F. Javed, H. Md. Akil, M. Jadoon Khan, M. Safdar, I. U. Din, M. A. Alotaibi, A. I. Alharthi, M. A. Bakht, A. Ahmad, A. A. Nassar, International Journal of Biological Macromolecules 2023, 232, 123476.

[33] A. K. Miri, D. Nieto, L. Iglesias, H. Goodarzi Hosseinabadi, S. Maharjan, G. U. Ruiz-Esparza, P. Khoshakhlagh, A. Manbachi, M. R. Dokmeci, S. Chen, S. R. Shin, Y. S. Zhang, A. Khademhosseini, Advanced Materials 2018, 30, 1800242.

[34] I. Pepelanova, K. Kruppa, T. Scheper, A. Lavrentieva, Bioengineering 2018, 5, 55.

[35] M. Kesti, P. Fisch, M. Pensalfini, E. Mazza, M. Zenobi-Wong, BioNanoMaterials 2016, 17, 193.

[36] D. R. Griffin, W. M. Weaver, P. O. Scumpia, D. Di Carlo, T. Segura, Nature Mater 2015, 14, 737.

[37] J. Madrid-Wolff, A. Boniface, D. Loterie, P. Delrot, C. Moser, Advanced Science 2022, 9, 2105144.

[38] H. Liu, P. Chansoria, P. Delrot, E. Angelidakis, R. Rizzo, D. Rütsche, L. A. Applegate, D. Loterie, M. Zenobi-Wong, Advanced Materials 2022, 34, 2204301.

[39] R. C. Hurley, S. A. Hall, J. E. Andrade, J. Wright, Phys. Rev. Lett. 2016, 117, 098005.

[40] X. Li, S. Chen, J. Li, X. Wang, J. Zhang, N. Kawazoe, G. Chen, Polymers 2016, 8, 269.

[41] H. E. (박희언) Park, N. Gasek, J. (황정욱) Hwang, D. J. Weiss, P. C. (이) Lee, Physics of Fluids 2020, 32, 033102.

[42] W. Schuurman, P. A. Levett, M. W. Pot, P. R. van Weeren, W. J. A. Dhert, D. W. Hutmacher, F. P. W. Melchels, T. J. Klein, J. Malda, Macromolecular Bioscience 2013, 13, 551.

[43] L. Rebers, T. Granse, G. E. M. Tovar, A. Southan, K. Borchers, Gels 2019, 5, 4.

[44] A. C. Daly, M. D. Davidson, J. A. Burdick, Nat Commun 2021, 12, 753.

[45] D. Loterie, P. Delrot, C. Moser, Nat Commun 2020, 11, 852.

[46] R. Rizzo, D. Ruetsche, H. Liu, M. Zenobi-Wong, Advanced Materials 2021, 33, 2102900.

[47] R. J. H. Stenekes, O. Franssen, E. M. G. van Bommel, D. J. A. Crommelin, W. E. Hennink, Pharm Res 1998, 15, 557.

[47] A. Lee, A. R. Hudson, D. J. Shiwarski, J. W. Tashman, T. J. Hinton, S. Yerneni, J. M. Bliley, P. G. Campbell, A. W. Feinberg, Science 2019, 365, 482.

[49] J. Malda, J. Visser, F. P. Melchels, T. Jüngst, W. E. Hennink, W. J. A. Dhert, J. Groll, D. W. Hutmacher, Advanced Materials 2013, 25, 5011.

[50] L. R. Nih, E. Sideris, S. T. Carmichael, T. Segura, Advanced Materials 2017, 29, 1606471.

[51] D. Lam, H. A. Enright, J. Cadena, S. K. G. Peters, A. P. Sales, J. J. Osburn, D. A. Soscia, K. S. Kulp, E. K. Wheeler, N. O. Fischer, Sci Rep 2019, 9, 4159.

[52] J. Kajtez, M. F. Wesseler, M. Birtele, F. R. Khorasgani, D. Rylander Ottosson, A. Heiskanen, T. Kamperman, J. Leijten, A. Martínez-Serrano, N. B. Larsen, T. E. Angelini, M. Parmar, J. U. Lind, J. Emnéus, Advanced Science 2022, 9, 2201392.

[53] E. Sideris, D. R. Griffin, Y. Ding, S. Li, W. M. Weaver, D. Di Carlo, T. Hsiai, T. Segura, ACS Biomater. Sci. Eng. 2016, 2, 2034.

[54] M. N. Collins, C. Birkinshaw, Carbohydrate Polymers 2013, 92, 1262.

[55] N. Alkhouli, J. Mansfield, E. Green, J. Bell, B. Knight, N. Liversedge, J. C. Tham, R. Welbourn, A. C. Shore, K. Kos, C. P. Winlove, American Journal of Physiology-Endocrinology and Metabolism 2013, 305, E1427.

[56] Z. Sun, B. D. Gepner, S.-H. Lee, J. Rigby, P. S. Cottler, J. J. Hallman, J. R. Kerrigan, Acta Biomaterialia 2021, 129, 188.

[57] L. Dilworth, A. Facey, F. Omoruyi, International Journal of Molecular Sciences 2021, 22, 7644.

[58] T. Ronti, G. Lupattelli, E. Mannarino, Clinical Endocrinology 2006, 64, 355.

[59] P. S. Gungor-Ozkerim, I. Inci, Y. S. Zhang, A. Khademhosseini, M. R. Dokmeci, Biomater. Sci. 2018, 6, 915.

[60] J. Adhikari, A. Roy, A. Das, M. Ghosh, S. Thomas, A. Sinha, J. Kim, P. Saha, Macromolecular Bioscience 2021, 21, 2000179.

[61] O. Jeon, Y. B. Lee, H. Jeong, S. J. Lee, D. Wells, E. Alsberg, Mater. Horiz. 2019, 6, 1625.

[62] D. Wu, Y. Yu, J. Tan, L. Huang, B. Luo, L. Lu, C. Zhou, Materials & Design 2018, 160, 486.

[63] D. Ribezzi, R. Pinos, L. Bonetti, M. Cellani, F. Barbaglio, C. Scielzo, S. Farè, Frontiers in Biomaterials Science 2023, 2.

[64] P. Datta, M. Dey, Z. Ataie, D. Unutmaz, I. T. Ozbolat, npj Precis. Onc. 2020, 4, 1.

[65] E. M. Langer, B. L. Allen-Petersen, S. M. King, N. D. Kendsersky, M. A. Turnidge, G. M. Kuziel, R. Riggers, R. Samatham, T. S. Amery, S. L. Jacques, B. C. Sheppard, J. E. Korkola, J. L. Muschler, G. Thibault, Y. H. Chang, J. W. Gray, S. C. Presnell, D. G. Nguyen, R. C. Sears, Cell Reports 2019, 26, 608.

[66] A. Ribeiro, M. M. Blokzijl, R. Levato, C. W. Visser, M. Castilho, W. E. Hennink, T. Vermonden, J. Malda, Biofabrication 2017, 10, 014102.

[67] V. H. M. Mouser, F. P. W. Melchels, J. Visser, W. J. A. Dhert, D. Gawlitta, J. Malda, Biofabrication 2016, 8, 035003.

[68] S. Chen, W. S. Tan, M. A. Bin Juhari, Q. Shi, X. S. Cheng, W. L. Chan, J. Song, Biomed. Eng. Lett. 2020, 10, 453.

[69] T. J. Hinton, A. Hudson, K. Pusch, A. Lee, A. W. Feinberg, ACS Biomater. Sci. Eng. 2016, 2, 1781.

[70] M. Becker, M. Gurian, M. Schot, J. Leijten, Advanced Science n.d., n/a, 2204609.

[71] P. N. Bernal, P. Delrot, D. Loterie, Y. Li, J. Malda, C. Moser, R. Levato, Adv. Mater. 2019, 31, 1904209.

[72] G. Größbacher, M. Bartolf-Kopp, C. Gergely, P. N. Bernal, S. Florczak, M. de Ruijter, J. Groll, J. Malda, T. Jüngst, R. Levato, Adv Mater 2023.doi: 10.1002/adma.202300756

[73] C. M. Rackson, K. M. Champley, J. T. Toombs, E. J. Fong, V. Bansal, H. K. Taylor, M. Shusteff, R. R. McLeod, Additive Manufacturing 2021, 48, 102367.

[74] B. E. Kelly, I. Bhattacharya, H. Heidari, M. Shusteff, C. M. Spadaccini, H. K. Taylor, Science 2019, 363, 1075.

[75] K. S. Lim, R. Levato, P. F. Costa, M. D. Castilho, C. R. Alcala-Orozco, K. M. A. van Dorenmalen, F. P. W. Melchels, D. Gawlitta, G. J. Hooper, J. Malda, T. B. F. Woodfield, Biofabrication 2018, 10, 034101.

[76] M. Mansouri, S. Xue, M.-D. Hussherr, T. Strittmatter, G. Camenisch, M. Fussenegger, Small 2021, 17, 2101939.

[77] A. Essaouiba, R. Jellali, M. Shinohara, B. Scheidecker, C. Legallais, Y. Sakai, E. Leclerc, Journal of Biotechnology 2021, 330, 45.

[78] D. Balboa, T. Barsby, V. Lithovius, J. Saarimäki-Vire, M. Omar-Hmeadi, O. Dyachok, H. Montaser, P.-E. Lund, M. Yang, H. Ibrahim, A. Näätänen, V. Chandra, H. Vihinen, E. Jokitalo, J. Kvist, J. Ustinov, A. I. Nieminen, E. Kuuluvainen, V. Hietakangas, P. Katajisto, J. Lau, P.-O. Carlsson, S. Barg, A. Tengholm, T. Otonkoski, Nat Biotechnol 2022, 40, 1042.

[79] A. Garcia, A. Sekowski, V. Subramanian, D. L. Brasaemle, Journal of Biological Chemistry 2003, 278, 625.

[80] C. Sztalryd, D. L. Brasaemle, Biochim Biophys Acta 2017, 1862, 1221.

[81] K. S. Lim, F. Abinzano, P. N. Bernal, A. Albillos Sanchez, P. Atienza-Roca, I. A. Otto, Q. C. Peiffer, M. Matsusaki, T. B. F. Woodfield, J. Malda, R. Levato, Advanced Healthcare Materials 2020, 9, 1901792.

[82] M. L. Terpstra, J. Li, A. Mensinga, M. de Ruijter, M. H. P. van Rijen, C. Androulidakis, C. Galiotis, I. Papantoniou, M. Matsusaki, J. Malda, R. Levato, Biofabrication 2022, 14, 034104.

[83] D. Scholz, D. Pöltl, A. Genewsky, M. Weng, T. Waldmann, S. Schildknecht, M. Leist, J Neurochem 2011, 119, 957.

[84] M. Imamura, E. Ozawa, Proc Natl Acad Sci U S A 1998, 95, 6139.

[85] E. P. Knott, M. Assi, D.D. Pearse, Biomed Res Int 2014, 2014, 651625.

[86] K. Krawczyk, S. Xue, P. Buchmann, G. Charpin-El-Hamri, P. Saxena, M.-D. Hussherr, J. Shao, H. Ye, M. Xie, M. Fussenegger, Science 2020, 368, 993.

